# A New Sensory Skill Shows Automaticity and Integration Features in Multisensory Interactions

**DOI:** 10.1101/2021.01.05.425430

**Authors:** James Negen, Laura-Ashleigh Bird, Heather Slater, Lore Thaler, Marko Nardini

## Abstract

People can learn new sensory skills that augment their perception, such as human echolocation. However, it is not clear to what extent these can become an integral part of the perceptual repertoire. Can they show automatic use, integrated with the other senses, or do they remain cognitively-demanding, cumbersome, and separate? Here, participants learned to judge distance using an echo-like auditory cue. We show that use of this new skill met three key criteria for automaticity and sensory integration: (1) enhancing the speed of perceptual decisions; (2) processing through a non-verbal route and (3) integration with vision in an efficient, Bayes-like manner. We also show some limits following short training: integration was less-than-optimal, and there was no mandatory fusion of signals. These results demonstrate key ways in which new sensory skills can become automatic and integrated, and suggest that sensory augmentation systems may have benefits beyond current applications for sensory loss.

## BODY

Sensory substitution and augmentation systems, such as human echolocation, seek to augment perception by providing an alternative way to infer information about the surrounding environment (Maidenbaum & Abboud, 2014). In practice, most of these devices and techniques have been attempts to replace vision for people who are blind. For example, the eye cane is a head-mounted device that translates distance measurements into vibrotactile signals, vibrating less when it is pointed at something further away (Maidenbaum et al., 2014). Human echolocation is a technique of using reflected mouth ‘clicks’ or other sounds to infer the spatial layout and material properties of objects in the surrounding environment (Kolarik et al., 2014; Thaler & Goodale, 2016). Taking advantage of a sensory substitution and augmentation system requires learning a new sensory skill. We define the term *new sensory skill* to mean a new ability to use information via an available sense. For example, learning to use an audio delay between emission and echo to judge distance.

Recent theory emphasizes broad scope to learn new sensory skills to a very high level, enabled by a remarkable degree of brain plasticity (Amedi et al., 2017; Auvray & Myin, 2009; Chebat et al., 2015; Striem-Amit et al., 2012). For example, expert echolocators can discriminate between a disc that is 50cm away versus 53cm away on 75% of trials (Thaler et al., 2019). These skill levels can be sufficient to allow everyday navigation, such as riding a bicycle through the woods (National Public Radio, 2011). These behavioural abilities are enabled by brain plasticity in regions that underlie perception, which are increasingly understood to represent information in a flexible amodal manner. For example, the ‘visual’ (occipital) cortex of expert echolocators responds to spatialized echo sounds (Thaler & Goodale, 2016) in a way that mimics retinotopic organization (Norman & Thaler, 2019). These and other findings (Amedi et al., 2017) suggest that the ‘visual’ cortex is actually an adaptive and amodal centre for processing detailed spatial information via whichever sense(s) can provide it. This background of high-level skills leads to an exciting scope for further research involving new sensory skills in more diverse settings.

While previous results definitively demonstrate that people can learn to use new sensory skills to a high level, it remains unclear to what extent these skills can become automatic and integrated in a similar way to the rest of perception. One possibility is that new sensory skills, particularly those learned in adulthood, will remain slow, perhaps requiring explicit linguistic support, and separated from other perceptual information. It has been suggested that the use of an augmented system may remain fundamentally different from native perception (Deroy & Auvray, 2012). Computational studies have also suggested that learning to integrate estimates from a new sensory skill into multisensory processing should take hundreds of thousands of exposures (Daee et al., 2014), making it very impractical to learn. On the other hand, findings of radical reconfiguration of sensory processing in some populations who have learned alternative sensory skills show the great potential flexibility of human perceptual mechanisms (Amedi et al., 2017).

One major key to determining when perception shows automatic and integrated features is a shift to multisensory paradigms. Most studies regarding sensory augmentation focus on unisensory tasks (where the participant can only use the new sensory skill), so there is a major gap in our knowledge of how these new sensory skills operate during multisensory interactions. This is important because in everyday perception, multiple modalities are integrated to build the overall experience and interpretation of the surrounding world. For example, vision and sound are combined to localise objects (the ventriloquist effect; Alais & Burr, 2004), and brief sounds and flashes are combined to count events (the sound-induced flash illusion; Shams et al., 2002). When signals from different modalities are combined to drive the overall percept, this combination of estimates provides some key advantages such as the use of Bayesian principles to reduce sensory uncertainty (Ernst & Banks, 2002; Maloney & Mamassian, 2009). To understand how a new sensory skill participates in perception - particularly, the extent to which it is integrated alongside the other senses - it is crucial to study how it operates in multisensory tasks.

Our broad approach to understanding the potential of new sensory skills is to examine how new sensory skills function in terms of key multisensory interactions, using known results regarding native perception as a guide for creating hypotheses. For example, we test whether new signals can enhance the speed of decision-making. Scientifically, this approach seeks to extend our knowledge of how flexibly perception is organised in the brain (Amedi et al., 2017). Practically, it seeks to extend the potential user base for new sensory skills, because a new sensory skill that is integrated into perception (e.g. participating in efficiently integrated sensory computations) could be useful to a much broader user base than a new sensory skill that is isolated. This user base might not only include people who need to replace vision, but also people who want to enhance their perception in other ways. For example, a surgeon who uses sound to guide their operation or a person with moderate vision loss who wants to play sports with the aid of new technologies. Please note that we are not making any comment on the scope for a “new sense”, the subjective experience of sensory substitution (König et al., 2016), or the possibility of creating “visual” experiences through non-visual stimuli (Abboud et al., 2014; Deroy & Auvray, 2012). Instead, our specific aim is to understand the principles by which new sensory skills function in multisensory environments; our approach is to compare new and native perceptual skills in functional terms – in particular: automaticity and cue combination.

As a testbed, this study uses a new sensory skill inspired by human echolocation (Kolarik et al., 2014; Thaler & Goodale, 2016). Participants hear two identical clicks in series. A longer delay between two auditory clicks indicates that the target is further away. After guessing, participants have visual feedback of the target’s true distance. This is less complex than real echolocation, because it does not involve any change in amplitude or power spectrum between the emission and the echo, the potential for different materials or shapes to reflect sounds differently, nor any variation in the emission itself (Zhang et al., 2017). However, it makes for a tightly controlled model system, in which participants must learn to map a single cue (auditory delay) to a single parameter (distance). It also parallels the case of a device for sensory substitution/augmentation that translates a single physical property to a sensory cue – for example, the EyeCane (Maidenbaum et al., 2014) that translates distance to auditory pitch and/or vibration intensity. In our study, participants are trained to judge distances with a new auditory delay cue. This training backdrop is used to explore multisensory functions (specifically: automaticity and cue combination) in the new sensory skill.

### Automaticity

Fully defining what it means for a sensory skill to be “automatic” is difficult, but it involves features such as speed, non-verbal processing, lack of explicit control, a subjective experience of lacking effort, and so on (Moors & De Houwer, 2006). In other words, automatic sensory-motor processing does not require much deliberation or verbalisation, so that it can take place while we also do something else – such as walking or driving while holding a conversation. Fully testing for automaticity is not feasible, so this report focuses on automaticity features that are particularly relevant to multisensory processing and the potential of sensory augmentation approaches.

One of the automaticity features we focus on is non-verbal processing of the stimulus. For example, non-verbal processing for echolocation would mean that you could simply hear how far away things are based on their echoes without any language or explicit reasoning. Verbal processing, on the other hand, would mean that you covertly used language (‘in your head’) to reason about the distance of an object based on sounds that you heard. This verbalization might for example entail a reminder that a longer delay between emission and echo indicates a further target and the use of language to reason about what you heard. It might take a form of a statement, such as “The second delay was longer. That target must have been further.” Importantly, non-verbal processing leaves verbal resources free for other uses. To address this, we tested for non-verbal processing using a dual-task paradigm; we asked participants to judge echoic distances in the presence or absence of a simultaneous verbal working memory task. If people process echo-like cues non-verbally, the simultaneous task should not interfere with the ability to judge distance.

The second automaticity feature we focus on is speed, i.e. that processing should be ‘fast’. This is important because of its relevance to the everyday use of a new sensory skill in naturalistic settings. How ‘fast’ a skill should be is difficult to define. Thus, we decided to break this into two parts. In part one, we examined average reaction times descriptively, i.e. how fast are they? In part two, we examine a grounded benchmark of being ‘fast’ by looking for the so-called redundant signals effect (Hershenson, 1962; Miller, 1982). The redundant signals effect is observed when two cues (e.g. a visual and an auditory cue) to a simple choice or response (e.g. distance) lead to faster reaction times when presented together than the decision being made using either single cue alone. Therefore, the presence of a redundant signals effect indicates that a novel audio cue is processed on a similar (‘fast’) time scale as a visual cue. The absence of a redundant signals effect on the other hand signifies that the two cues are processed on vastly different timescales. For example, consider that processing of an ordinary visual cue took between 500 and 1000ms, whilst processing of a novel audio cue took between 5000 and 10000ms. In that case, we would not see a redundant signals effect because a simultaneous novel audio cue could not improve the speed offered by the ordinary visual cue. Therefore, the presence of a redundant signals effect indicates that a novel audio cue is processed on an overlapping (‘fast’) time scale as a visual cue. This will serve as our second working definition of ‘fast’. Please note that we will not be addressing the extensive debate on ‘racing’ or ‘pooling’ / ‘coactivation’ (Otto & Mamassian, 2017). We are just trying to see if the novel audio cue can be processed fast enough to have any benefit on top of a more ordinary visual cue. (As it happens, any redundant signals effect will also speak somewhat to the issue of using multiple cues efficiently, in line with the questions below related to cue combination.)

The third and final automaticity feature we focus on is forced fusion (Hillis et al., 2002). Forced fusion is an indicator of automaticity because participants subject to fusion “cannot help but” combine sensory estimates, even though when it makes their performance worse; in other words, this aspect of perception is “goal independent” (Moors & De Houwer, 2006). Forced fusion is when two separate estimates of the same feature of the world are combined and the participant loses access to the two pre-combination estimates in favour of the single post-combination estimate. For example, when you look at a tabletop, it is relatively easy to judge the tabletop’s slant (usually 90 degrees from vertical). However, judgments of slant (and other aspects of spatial layout) come from a range of cues such as binocular disparity, texture, linear perspective, motion parallax, and so on. Although these visual cues may lead to slightly differing estimates of slant, we are unable to perceive them separately from each other. One potential advantage of forced fusion is that perception can be improved by an accurate prior expectation that some signals will be strongly correlated (Ernst, 2006). Following standards in the field, we will test for forced fusion with an oddball task (Hillis et al., 2002).

### Cue combination

We also looked for integration by testing for the combination of different cues in a Bayes-like manner to reduce noise (uncertainty). This effect has been found many times in native multisensory perception (Alais & Burr, 2004; Ernst & Banks, 2002; Knill & Pouget, 2004; Pouget et al., 2013). Suppose that both vision and audition signal something about the state of the world, like the location of a speaker. If both senses have Gaussian noise around the correct target, then the lowest possible noise can be achieved by taking a reliability-weighted average of the best estimates from each system (Alais & Burr, 2004). Specifically, it becomes possible to achieve additive precision: with precision defined as 1/variance, with optimally-chosen weights, the combined estimate can have a precision that is the sum of the two individual precisions (Ernst & Banks, 2002). This is sometimes a major reduction in noise. A full mathematical treatment is discussed here (Ernst et al., 2016). Even when this is not implemented optimally, it can still reduce noise given two sensory modalities than either single modality alone. This is potentially an important advantage for difficult judgments or ones where precision is important. We will refer to this process as *cue combination*. Key evidence for cue combination is that responses are less noisy given two different cues together than either single cue alone. One major goal of the present study is to further examine if a new sensory skill can be combined with native perception to achieve a cue combination effect.

Previous results regarding cue combination with new sensory skills are mixed (Goeke et al., 2016; Negen et al., 2018) and one open question is if any discrepancy in previous results could be explained by a narrower issue: whether the sensory noise (uncertainty) is internal or external (Macmillan & Creelman, 2004). A cue with *internal noise* could be perfectly reliable, but its perception by humans remains noisy because of noise that occurs internally during perception. (Or at least, it could be so close to perfectly reliable that the difference is negligible.) A cue with external noise is one that is somewhat unreliable because of noise in how it is generated in the environment, even if the information it provides is perceived perfectly. In other words, there is noise that is external to perception. Since many everyday tasks involve significant levels of internal noise, it is important to establish if it is possible to achieve a cue combination effect with a native cue and a new sensory skill that are both subject to internal noise. We will do this by modelling aspects of the current study on the previous one that did find cue combination (Negen et al., 2018), except replacing the visual cue (which had external noise) with a visual cue that has internal noise. This should clarify the possibility of showing a cue combination effect with a new sensory skill and native perception even when there is only internal noise in the task.

### Study Overview

To resolve these research questions we had a group of participants go through training and testing with a new sensory skill over 10 sessions. We modelled the skill on human echolocation: the length of a delay between two auditory clicks maps linearly to a target’s distance from the observer. Participants were trained to judge distances with this new cue. We also introduced a simple visual cue to distance that is easy to understand but also subject to internal noise in perception. Participants were asked to judge distances with the audio cue, the visual cue, or both. We then tested for automaticity and cue combination by introducing specific variants to the basic task.

### Hypotheses

The study was designed to test a series of hypotheses organized into two themes: the ‘automaticity’ theme and the ‘cue combination’ theme. The automaticity theme tests predictions about whether participants’ use of the new audio cue will acquire some key ‘automatic’ properties during multisensory interactions. The ‘cue combination’ theme tests predictions about whether participants will combine a novel audio cue with a native visual cue in a Bayes-like manner even when that visual cue is subject to purely internal noise. Together, these tests will start to sketch out to what degree a new cue can become more like the perception involved when you look at a scene yourself (not requiring language, fast, fused, integrated) and less like having to reconstruct a scene from a verbal description (requires language, slow, unfused, singular). Each of these themes contains several specific hypotheses.

#### Cue Learning Hypothesis (Preliminary)

This hypothesis states that participants will learn how to use the novel audio cue on its own to make distance judgements. Previous research suggests that the use of this cue is very poor without training – worse than a simple strategy of pointing to the center of the line on each trial (Negen et al., 2018). Learning to use the novel audio cue is an essential first hurdle for participants to pass in order to test further hypotheses about how they use the new skill, i.e. automaticity and cue combination. We ask whether each participant learned the new cue by testing for a significant correlation between target distances and participants’ responses in the second training session.

#### Parallel to Verbal Hypothesis (Automaticity)

This hypothesis states that performance on the distance judgement task is not dependent on verbal or linguistic resources. In lay terms, this means that participants can use the echo-like cue without ‘talking themselves through it’, and so could potentially carry on other verbal tasks at the same time. We test this by comparing trials in which participants only judge distance, trials in which participants only recall a probe digit from a string of six verbal digits (verbal working memory task), and trials in which they do both at the same time. This hypothesis states that variable error in the judgements will be similar (i.e. not significantly different) with and without the verbal working memory task; and that the proportion of digits recalled correctly will be similar with and without the distance judgement task.

#### Redundant Signals Hypothesis (Automaticity)

Being automatic involves processing a cue ‘fast’, which is notoriously difficult to define. We will first report descriptive statistics for the reader to judge. We also choose to look for a redundant signals effect: giving participants two cues to a simple choice leads to faster (but not less accurate) choice responses than with either single cue alone. We test this by giving participants very easy distance judgments and ask them to prioritise making their choice quickly. We hypothesize that mean accuracy will be above chance for the audio cue alone; that mean reaction time will be lower with both cues than the best single cue alone; and that mean accuracy will be similar with both cues and the best single cue alone (i.e. gaining speed without sacrificing accuracy). For this to happen, the speed of processing of the novel audio cue must be competitive with the speed of processing of the native visual.

#### Forced Fusion Hypothesis (Automaticity)

This hypothesis states that the audio and visual cue will become subject to forced fusion. This means that participants will lose (some) access to the perception of distances indicated by each individual cue and instead only have access to their combined perception of the distance. We test this using an oddball task. Two choices are given that are exactly the same, with both an audio and visual signal indicating a standard distance. The third choice is the comparison. The three choices are presented in random order and the participant’s task is to indicate which choice was different from the other two. There are two sub-types. The first is the congruent comparison, where audio and visual components are either both further than the standard (A+V+) or both nearer than the standard (A-V-). This should be unaffected by forced fusion and serve as a baseline. The second is the incongruent comparison, where there is one cue indicating a further distance and the other indicating a nearer distance (A-V+ or A+V-). If forced fusion is occurring, these should be perceived as very similar to the standard because the opposing deviations should be cancelled out when they are fused into a single distance percept (Hillis et al., 2002). If this hypothesis is correct, then incongruent performance should be worse than congruent performance.

#### Precision Hypothesis (Cue Combination)

This hypothesis states that participants will combine the two cues to gain precision (reduce noise). This is a key efficient multisensory interaction that flexibly uses the senses to provide reasonable accuracy even when specific senses fail to provide reliable estimates. We test for cue combination by asking whether variable error (the part of error that is just noise rather than systematic distortion) is lower when both cues are available than when the best single cue is available (Ernst et al., 2016). A finding of reduced variable error (cue combination) would replicate previous work (Negen et al., 2018) but with a cue that has internal noise.

#### Optimal Weight Hypothesis (Cue Combination)

The above hypothesis suggests that people will combine cues to gain some precision. This hypothesis extends this further: that people will gain near-optimal levels of precision by setting the relative weight they give to each cue near-optimally, according to its relative reliability. This hypothesis has four parts: the actual weight participants give to each cue will be correlated with the optimal weight; the average weight given to each cue will be similar to (not significantly different from) the average optimal weight; average weights given to the visual cue will increase when the visual cue is made more reliable; the variable error with both cues will be similar to the optimal variable error that is theoretically possible. These are a standard set of criteria for testing whether near-optimal Bayesian cue combination is occurring (Ernst et al., 2016).

## Methods Overview

In the *Full Methods* section, we give the full details required to replicate the study. Because this information is extensive and detailed, we first summarize it here with the goal of clarifying the intent and key parts of the approach.

Throughout the study, a virtual cartoon whale named Patchy was presented in an immersive 3D environment and gave instructions, encouragement, and feedback to participants (see Figure 1, right panel). Judging the distance to patchy using one or more sensory cues was the main task on each trial. He could hide under the virtual sea along a line that stretched out in front of the participant, who was sitting on a chair at the very front of a boat. As a baseline, it is helpful to look at an AV trial (audio-visual; Figure 1). The participant is looking at the line in front of the boat. The goal is to estimate the distance to the non-visible target (Patchy). At the same time, they hear a pair of clicks (auditory cue) and see a cluster of blocks distributed along the line (visual cue). The auditory cue is informative about Patchy’s position because the delay between the first click and a second click is proportionate to distance (like an echo). The visual cue is informative about Patchy’s position because the blocks move (contract) towards a convergence point which is Patchy’s location (see Figure 1). During 250ms (overlapping with the audio delay), the blocks move 20% of the way towards the target, then disappear. The participant uses an Xbox controller to control a marker which they use to indicate their estimate of Patchy’s distance along the line. When they have moved the marker to the correct location, they select the location by pressing a button. Patchy then surfaces at the correct location, serving as visual feedback of the correct location. He also gives feedback in the form of a percentage expressing their degree of under / overestimation. This completes an AV trial. In summary, we first trained participants to estimate distance using only the novel auditory stimuli. We then introduced the visual stimuli to the trials. Finally, we carried out several variations on these basic trial types to test specific hypotheses.

**Figure 1:**
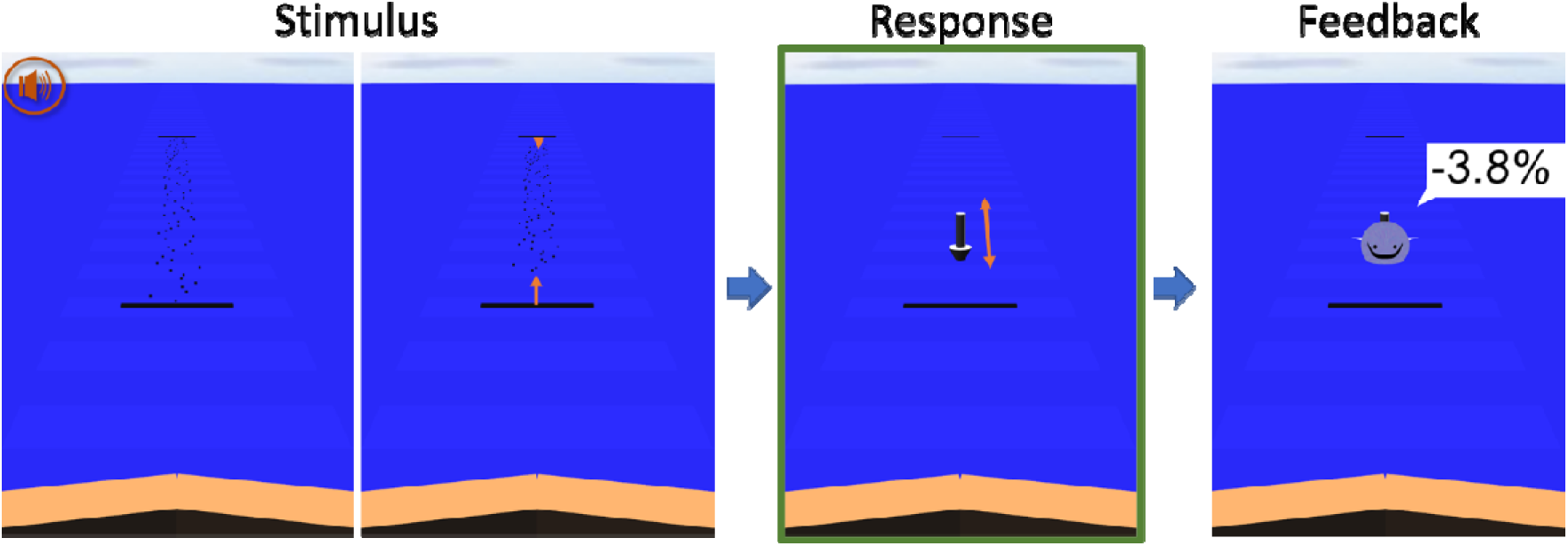
Example AV (audio-visual) trial. Everything in orange here (audio icon and orange arrows) was not seen by the participant and is edited on top for the reader’s clarity. Looking out over the front of the virtual boat, the participant sees two black bars that mark the range of locations in which the target fall. They hear an audio stimulus, two clicks in series where the delay signals distance. They also see a distribution of black blocks. The blocks all move 20% of the way towards the target over 250ms. The participant then controls a grey 3D marker which they can move between the black bars. When they press A and enter a response, Patchy appears at the correct location and provides feedback in terms of percent error. To be as clear as possible: the black bars, the black blocks, the gradient in the sea, the grey 3D marker, the cartoon whale, and the cartoon whale’s speech bubble are all seen by the participant in stereoscopic 3D.

The experiment involved a total of ten sessions, each lasting around 1 hour and carried out over a 2- to 10-week period. The first two sessions gave participants practice with the audio cue and the visual cue separately. Training followed a scaffolding approach, in line with our previous learning paradigm (Negen et al., 2018). Specifically, training with the audio cue began with initial trials made as simple as possible: 2-alternative forced-choice judgments for a 10m vs 35m target distance. The number of options was then increased via a 3- and then 5-alternative forced-choice design. This progressed to continuous judgement trials in which participants can use the joystick to respond anywhere along the line. Continuous responses in the second session are analysed to test the Cue Learning Hypothesis (Preliminary).

In session 3, AV trials were introduced. These were accompanied by Audio trials and Visual trials (like AV but with only one cue). This allows for a test of the Precision Hypothesis (Cue Combination). Session 4 repeated session 3. From session 5 onwards, there were variations to test more hypotheses. Session 5 asked participants to do Dual Task trials, in which they had to do an Audio/Visual/AV trial while remembering a string of six verbal digits. This was for the Parallel to Verbal Hypothesis (Automaticity). Sessions 6 and 7 had perturbation trials, in which the audio and visual cue were offset by 10%. This allowed us to measure the relative weight given to each cue (through multiple regression) and test the Optimal Weight Hypothesis (Cue Combination). Session 8 was the oddball task (Figure 2, top row), in which participants were shown the stimuli for 3 AV trials and asked to indicate which one was different from the other two. This was for the Forced Fusion Hypothesis (Automaticity), which specifically predicts that performance will be better when the oddball stimulus is congruent than incongruent. Session 9 was a speeded task (Figure 2, bottom row). Instead of making a slow and careful judgement, participants were given two possible distances and asked to rapidly indicate which of the two was correct. This tested the Redundant Signals Hypothesis (Automaticity). Finally, in session 10, there were dual task control trials. In these, participants completed the verbal working memory task without having to judge any distances. This was done to further test the Parallel to Verbal Hypothesis (Automaticity).

**Figure 2:**
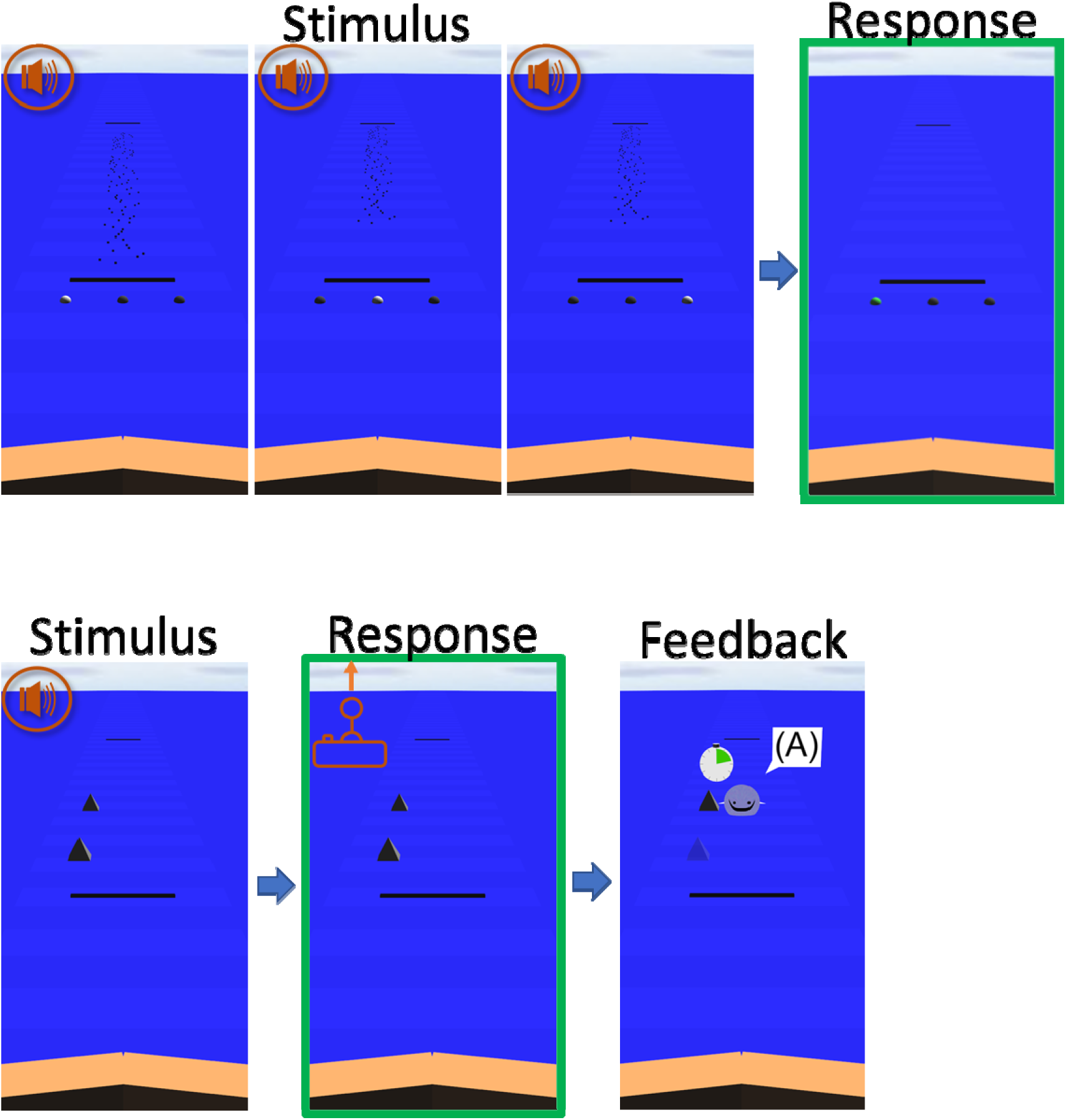
Oddball (top) and Speed Task (bottom) example trials. Everything in orange is drawn on top of the screenshots for the clarity of the reader (audio icon, joystick icon). For oddball, the participant is presented with three sets of AV stimuli, each matched with a sphere changing from black to white. Two stimuli are the same and one is different. The task is to indicate which AV stimulus was different (here, the first stimulus) by highlighting the sphere corresponding to the stimulus with the controller (here, the left sphere). There is no feedback. For the speeded task, the participant is given two possible choices. As fast as possible after the stimulus (trial shown here is audio-only, but there were also visual and AV), the participant pushes the joystick up or down to indicate the nearer or further option. If the choice is correct, Patchy appears and shows them their reaction time on a 3-second clock.

## Results

### Cue Learning (Preliminary) Hypothesis

This hypothesis was an essential first check that participants individually learned to use the new cue. Results were consistent with the hypothesis: of the fourteen participants, twelve showed evidence (significant correlations) for learning the audio cue in the second session. For those twelve, correlations ranged from 0.55 to 0.91, all p-values below 1×10^-26^; the remaining two were excluded. A post-hoc test for a practice effect was also run by entering the audio-only variable error for sessions 3 to 7 in a repeated measures ANOVA. This was not significant, F(4, 40) = 1.28, p = 0.294, suggesting that audio-only performance had largely stabilized by the end of the second session. Weber fractions ranged from 0.16 to 0.52 (mean of 0.28, median of 0.27, SD of 0.10). In other words, while performance was well above chance, participants were still in the early stages of learning and well short of the performance levels that experts achieve with their own clicks and non-virtual stimuli.

### Parallel to Verbal (Automaticity) Hypothesis

This hypothesis suggests that the new sensory skill can be used with minimal interference from a verbal work memory task. Results were consistent with the Parallel to Verbal (Automaticity) hypothesis. We found no evidence that the verbal working memory task interfered with the judgement task. There was no main effect of judgement-only (i.e. Audio, Visual or AV; sessions 3 and 4) versus Dual Task (session 5) on variable error, F(1,11) = 4.53, p = 0.057 (Figure 3, left). If anything, performance was trending towards better performance (lower variable error) with the dual task than without. Further, we found no evidence that the distance judgement task interfered with the verbal working memory task. The mean proportion of digits correctly recalled was not significantly different for the dual task trials versus the digit span control task (where there was no judgement to make), t(10) =-2.15, p = 0.057 (Figure 3, right). This suggests that processing these distance cues was largely parallel to verbal working memory.

**Figure 3:**
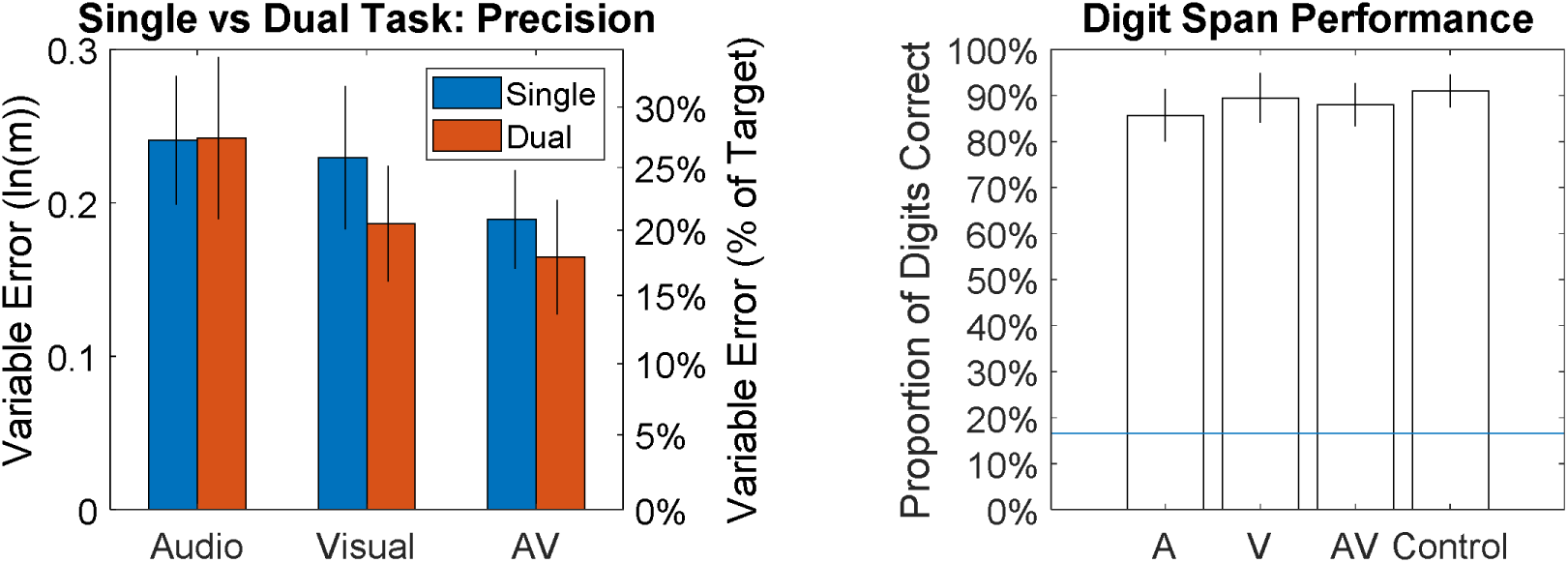
Performance for the dual task versus single-task variants. The left side charts variable error for different trial types. The right side charts the proportion of correctly answered memory probes. The blue line represents chance performance.

Further post-hoc testing also showed that variable error in the Dual Task AV trials was lower than variable error in the best single-cue dual-task trials, t(11) = −3.10, p = .010. In other words, we found a cue combination effect during trials where participants also had to complete a verbal working memory task. This further clarifies that the cue combination effect is not dependent on having verbal resources available.

### Redundant Signals (Automaticity) Hypothesis

This hypothesis tested whether participants could improve speed with both cues versus the best single cue, which is only possible if the new sensory skill can be used with some measure of speed. Results were consistent with the Redundant Signals (Automaticity) hypothesis. For Speed Audio trials, the mean reaction time was 740ms (95% CI: 628 to 851ms). Accuracy was significantly higher than the 50% accuracy benchmark for the Speed Audio trials, t(10) = 9.83, p < .001, d = 2.96, mean of 72% (95% CI: 68% to 78%). For all 11 participants, the best (fastest) cue was the visual cue. Mean reaction times were faster in Speed AV trials than Speed Visual trials, t(10) = 5.11, p <.001, d = 1.54 (Figure 4, bottom left). Choices were not significantly less accurate in Speed AV trials than Speed Visual trials, t(10) = 0.71, p = 0.496, d = 0.21 (Figure 4, bottom right). In other words, we found an improvement in speed that was not due to a trade-off in accuracy. Compared to Speed AV trials, Speed Audio trials were both slower, t(10) = 7.92, p < .001, d = 2.39, and less accurate, t(10) = −4.45, p = 0.001, d = −1.34. The central finding is that RTs and accuracy are better in Speed AV trials than in Speed Visual trials, which indicates that the processing of the new sensory audio signal is on timescale comparable to the ordinary visual cue, and therefore that it is ‘fast’.

**Figure 4:**
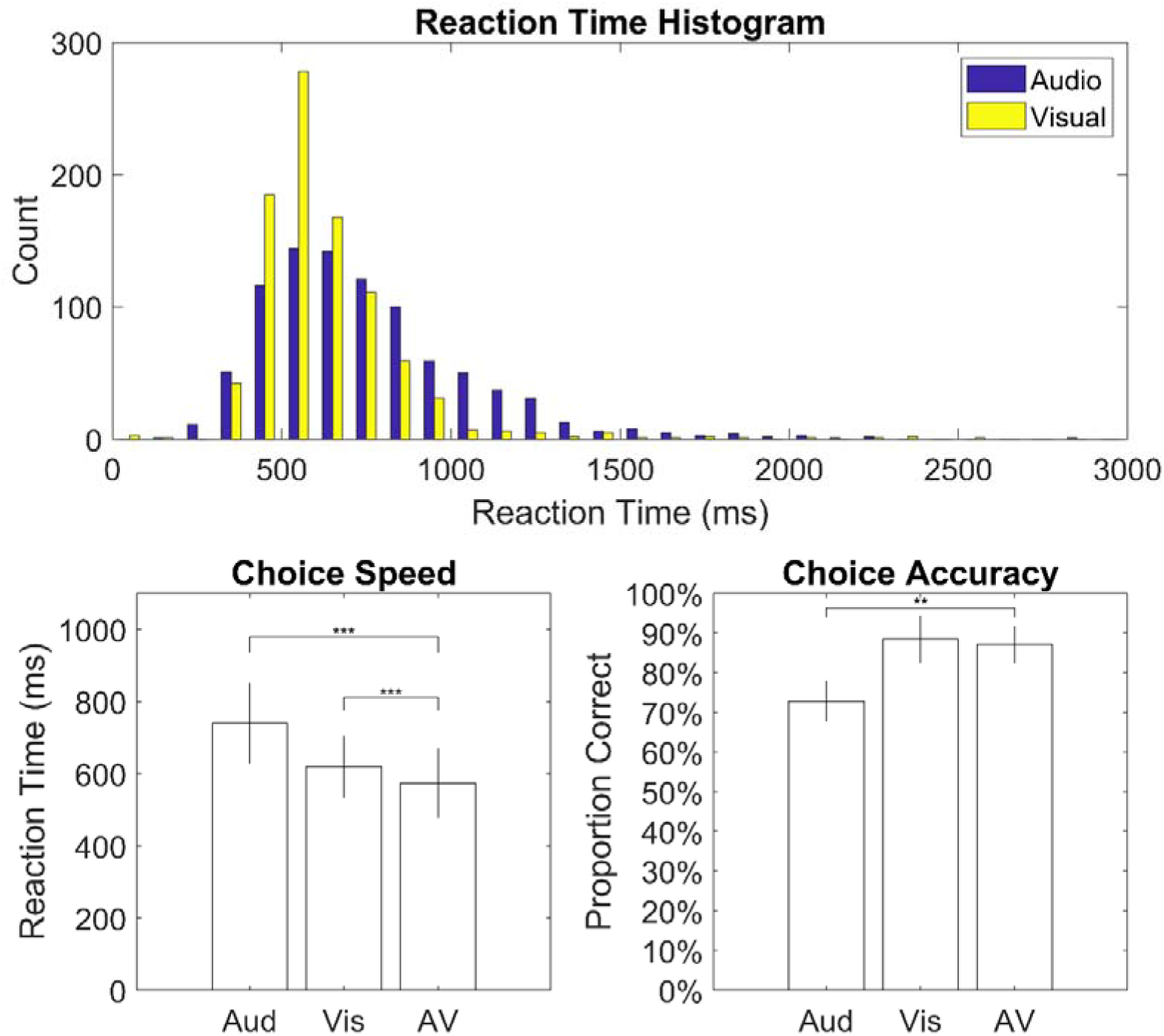
Performance on the speeded task by trial type. On the top, a histogram of reaction times to a choice of target versus target +/- 40%. On the bottom left, 90% trimmed means for reaction times. On the bottom right, accuracy of choices (50% chance). Participants were faster to make a choice with both stimuli (i.e. AV) than either single cue alone. Participants were also more accurate with both stimuli than with the audio cue alone.

### Forced Fusion (Automaticity) Hypothesis

This hypothesis suggests that participants will not be able to avoid averaging the two cues and thus perform worse on incongruent trials than congruent trials. We did not find any evidence in favour of the Forced Fusion (Automaticity) hypothesis. Fusion thresholds were not significantly different for congruent (A+V+ and A-V-) versus incongruent (A+V- and A-V+) oddball trial types, t(10) = 0.30, p = 0.768 for the mean deviation measure, t(10) = 1.32, p = 0.217 for the fitted threshold measure (Figure 5, left). We would expect performance to be worse in the incongruent trials if forced fusion was occurring. However, congruent and incongruent scores were highly correlated, r(9) = 0.92, p < .001 for the mean deviation measure, r(9) = 0.98, p < .001 for the fitted threshold measure (Figure 5, right). This suggests that the measure was reasonably sensitive to individual variation in precision at judging distance.

**Figure 5:**
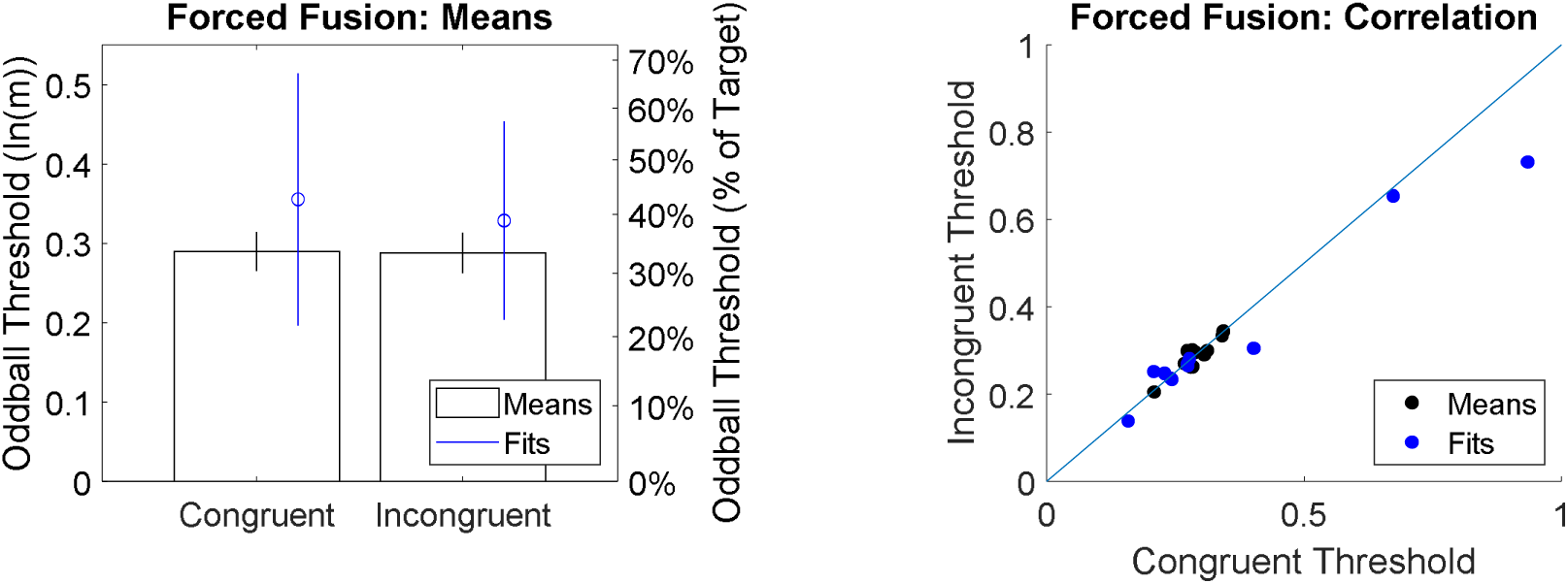
Results for the tests of forced fusion. On the left, thresholds were similar for congruent and incongruent trial types. Under forced fusion, we would expect incongruent performance to be worse. On the right, correlation between the two measures shows that the task was sensitive to individual variation in performance. Black dots and bars reflect estimating thresholds by the mean deviation. Blue dots and bars reflect estimating thresholds through a fitting method based on the normal distribution.

### Precision (Cue Combination) Hypothesis

This hypothesis tested whether participants gained precision (reduced variable error) by combining the novel auditory cue with a visual cue when both were available. Results are consistent with the Precision (Cue Combination) Hypothesis. The variable error for AV trials was lower (better) than the variable error for the best single cue, F(1, 11) = 5.59, p = .038, 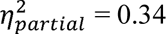. This was also true for a variation of the analysis where ranks were entered instead of raw scores, F(1, 11) = 5.92, p = .033, 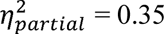. This suggests that participants did combine the new audio cue and native visual cue, extending previous results with external noise (Negen et al., 2018) to the present study using internal noise.

**Figure 6:**
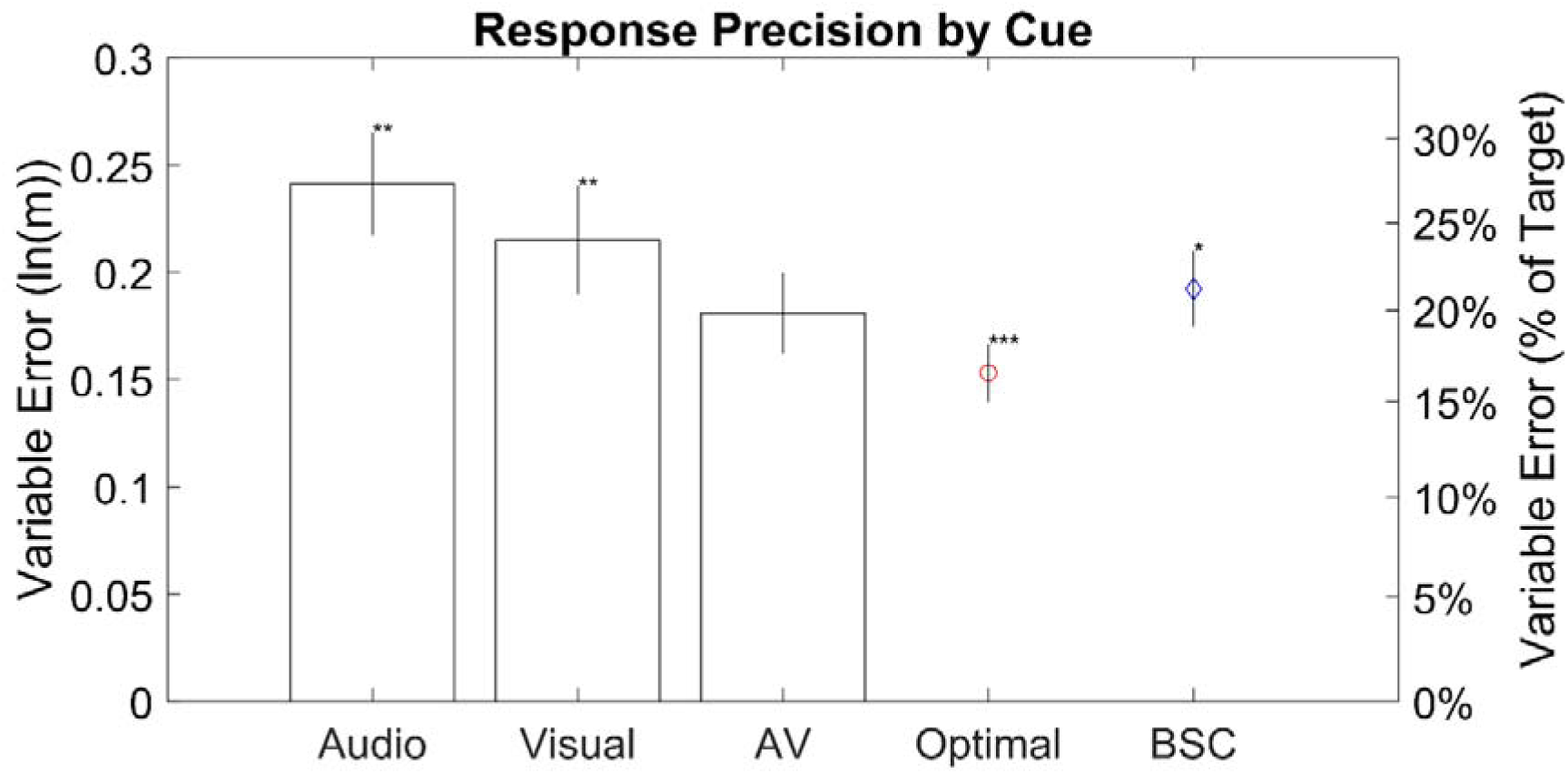
Average variable error by trial type in sessions 3 through 5. Variable errors are used to quantify the amount of non-systematic perceptual noise in judgements (see *Parallel to Verbal* under *Data Analysis* under *Method* for calculations). Audio and Visual are the trials where only the audio or only the visual stimulus, respectively, were presented. AV stands for audio-visual (i.e. trials where both the audio and visual stimulus were presented). Optimal and Best Single Cue (BSC) are not different trial types; they are values derived from the Audio, Visual, and AV trials. Optimal refers to the best possible variable error that should theoretically be possible with optimal computations. BSC stands for best single cue; for each participant and session, it is the lesser (better) of the audio variable error and the visual variable error. Error bars are 95% confidence intervals. Asterisks compare each type against AV in a paired t-test: *p < .05, **p < .01, and ***p < .001.

### Optimal Weight (Cue Combination) Hypothesis

This hypothesis tested whether participants set the optimal weights for each cue and gained optimal precision from the use of both cues. Results are partially consistent with the Optimal Weights (Cue Combination) Hypothesis. There was a significant correlation between optimal visual weights and observed visual weights, r(20) = 0.78, p < .001 (Fig 7). There was also a significant correlation between optimal audio weights and observed audio weights, r(20) = 0.80, p < .001 (Fig 7). Because the analysis procedure does not always result in observed weights that sum to exactly one (see Methods), both visual and auditory weights were separately analysed. When the visual reliability was higher in Session 7 than Session 6 (i.e. longer versus shorter), this did lead to higher weight being placed on the visual cue, t(10) = −3.15, p = 0.010, and lower weight being placed on the audio cue, t(10) = 3.93, p = 0.003 (Figure 7). Mean observed visual weight shifted from 0.588 for less reliable (i.e. shorter) visual cues to 0.776 for more reliable (i.e. longer) visual cues. All of this is consistent with the hypothesis. However, the observed visual weights were higher on average than the optimal visual weights, F(1, 10) = 5.53, p = 0.041. The mean observed visual weight was 0.682 versus an optimal 0.608. In addition, the observed audio weights were lower on average than the optimal audio weights, F(1, 10) = 16.32, p = 0.002. The mean observed audio weight was 0.259 versus an optimal 0.391. This suggests that participants did vary in how they set integration weights, and that this variation is explained partially by the correct optimal weight, but that participants also tended to systematically over-rely on the visual cue. In accordance with this, the variable error in AV trials was higher than the optimal variable error, F(1, 11) = 23.00, p < .001. This was also true in a variation where the ranks were entered instead of the raw variable errors, F(1, 11) = 32.69, p < .001.

**Figure 7:**
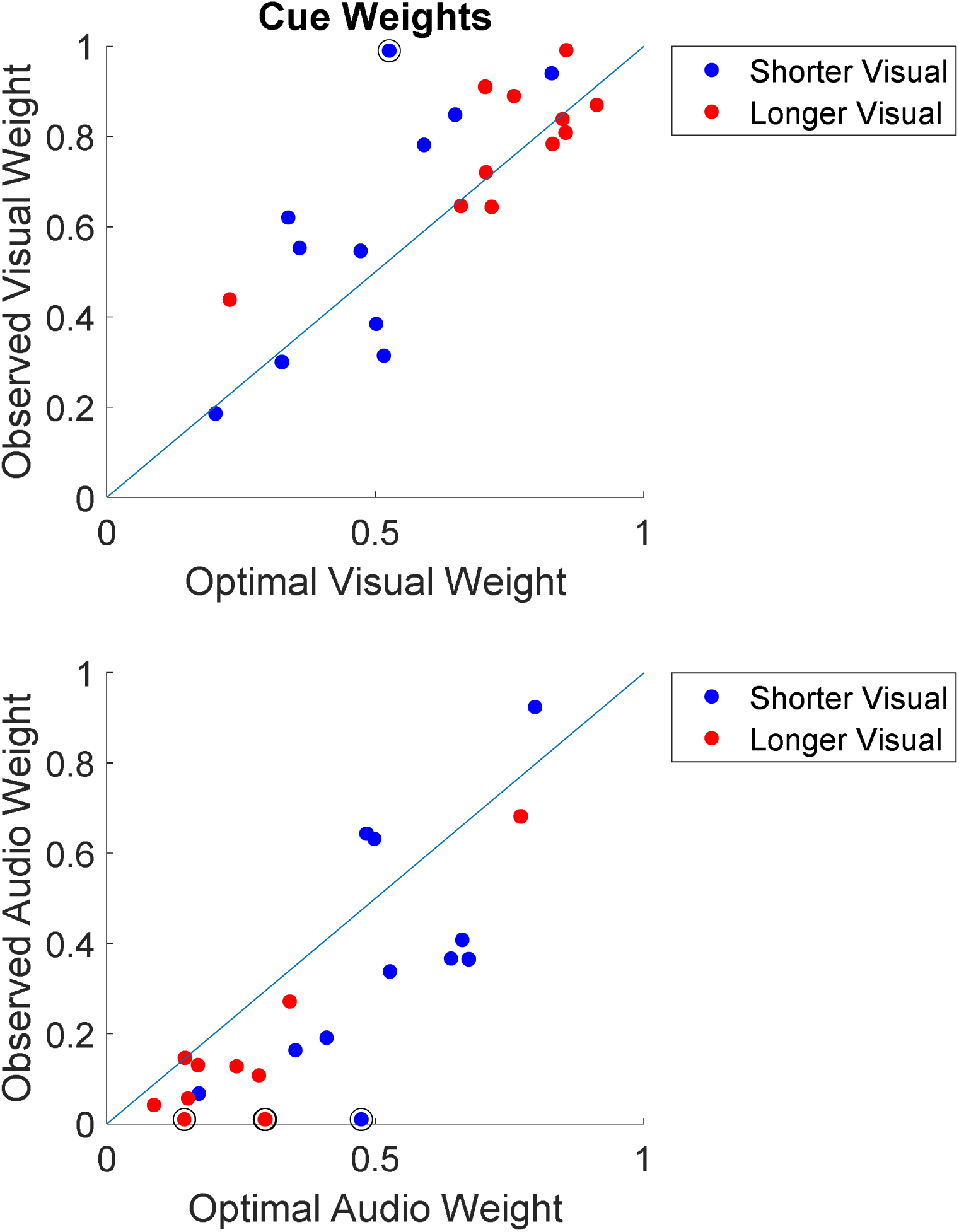
Weights given to each cue. Each blue dot is a separate participant in session 6, when the visual cue was shorter (less reliable). Each red dot is a separate participant in session 7, when the visual cue was longer (more reliable). Dots with a black circle around them were adjusted to be in the range of zero to one (i.e. into the range of prior plausibility). The blue line is the identity line.

Appendix A examines this interpretation via cross-validation. These data are used to compare three models: one where people combine cues optimally, one where people combine cues with too little weight on the new sensory skill, and one where participants only rely on one cue at a time. In summary, the model where people under-rely on the new sensory skill is favoured for eight of twelve participants, with the optimal model favoured for another three. This largely accords with the interpretation given here (Bayes-like cue combination, but with sub-optimal weights).

## Discussion

In this study, we wanted to understand the principles by which new sensory skills operate in multisensory environments, using native perception as a guide for generating hypotheses. We therefore looked for several effects that are frequently observed in native multisensory perception, but we here tested them with a new sensory skill (using audio delay to judge distance) and a simple native visual cue. We found that use of this new skill met three key criteria for automaticity and sensory integration: enhancing the speed of perceptual decisions, processing through a non-verbal route, and integration with vision in an efficient, Bayes-like manner. We also show limits: integration was less-than-optimal, and there was no mandatory fusion of signals. It is noteworthy that these skills were attained after only very short initial training and experience. Our results provide further evidence in keeping with proposals that plasticity in perception allows new sensory skills to take on functions similar to native perception (Amedi et al., 2017; Maidenbaum & Abboud, 2014; Striem-Amit et al., 2012).

One key theme centred around automaticity. Here we saw some encouraging and important results: processing of the new cue becomes non-verbal, or at least lacks major interference from verbal tasks; the new sensory skill is ‘fast’, meaning that it can assist vision in making speeded choices even faster (without losing accuracy). These suggest a significant scope for new sensory skills to be implemented in more ecologically valid tasks, since they can take up few verbal resources and can help with situations that require a rapid response. On the other hand, we did not see any evidence of forced fusion. Even for native perception, forced fusion is something of a trade-off; it reduces the overall load on memory and can improve precision under certain circumstances but it also sacrifices explicit access to sensory information. Our results suggest that a useful level of automaticity can be achieved quickly, but that some kinds of automaticity adaptations (like forced fusion) are either out of reach or may require different circumstances (for example, longer training).

Another key question was whether participants would show efficient cue combination with a cue that is subject to internal noise. We found a number of key findings consistent with this: responses were less noisy with both cues than either single cue alone; the weight given to each cue was positively correlated with the optimal weight, and the weight flexibly changed when cue reliabilities changed. In short, participants followed Bayes-like principles to gain measurable benefits from combining multiple cues, much as is seen in native multisensory perception (Alais & Burr, 2004; Ernst & Banks, 2002; Knill & Pouget, 2004; Pouget et al., 2013). This adds important knowledge to the new field of multisensory processing with new sensory skills – the only previous evidence for Bayes-like combination with a new sensory skill came from a study in which the other (visual) cue had external noise (Negen et al., 2018). Because internal noise is typically the major issue for real-world perceptual-motor problems, however, the current results link lab findings more closely to everyday perception and to potential future applications of sensory training.

However, while performance qualitatively followed Bayesian predictions, performance did not quantitatively meet the predictions of optimal noise reduction. The weight given to the native visual cue was too high; the weight given to the new sensory skill was too low; noise was not reduced to the optimal level theoretically possible. This suggests that new sensory skills can interact with native perception to reduce noise, but that optimal noise reduction is (again) either out of reach or may require different circumstances (for example, longer training). It should be noted however that full Bayesian optimality is a high standard to compare with, since even native perceptual skills do not always show optimal Bayesian performance (Rahnev & Denison, 2018), and full evaluations of optimality also require consideration of costs and priors, not included in this study.

Flexible cue combination is an especially important result because it expands the scope of sensory training from the traditional model of substitution or replacement to one of augmentation. In our study, the new signal did not only substitute for vision, but improved the precision of existing visual capabilities. This has important potential applications to patients with sensory loss (e.g. partial vision loss) who still have some useful visual function. It also has applications towards developing devices to further enhance healthy perception, such as the use of additional sensors to guide surgery, to navigate, to play enhanced sports, to more efficiently work with heavy objects in a warehouse, or to locate potential hazards.

If we expect that these results reflect ecological use of a sensory substitution or augmentation device/technique, then they give some reasons to think that such approaches hold a great deal more promise than their current applications. New sensory skills are not isolated in the stream of perceptual processing. They can be used to make careful decisions less noisy and make fast decisions even faster. They take up little space in verbal working memory. This is not necessarily going to be perfectly optimized (at least not in this kind of timescale), but it is enough that the new sensory skill is making a measurable contribution to the multisensory aspects of perception.

With all that said, we should clarify two limitations that may be important. First, there is no universal objective definition of a cognitive process being ‘fast’. While the new sensory skill met our definition of ‘fast’ here, it still took people an average of 740ms (95% CI: 628 to 851ms) to make a decision. Exactly how to judge a certain speed as adequate or inadequate will ultimately depend on the specific intended use.

Second, the present study was only designed to detect very large verbal interference effects. We consider this good evidence that performance with the new sensory skill does not, in layman’s terms, depend merely on ‘talking themselves through’ its use. However, it very much remains possible that a more extensive investigation of this issue will reveal smaller interference effects from verbal working memory. There may also be individual differences in terms of ability to process new sensory skills through non-verbal routes, which the present study was also not designed to detect.

The dual task results also bring up a very unexpected possibility for future work. If anything, participants performed better with the dual task. One participant spontaneously described this as such: “with all the numbers, you sort of get out of your head and just point to the right spot.” This is extremely speculative, but it may be that a distracting task can act as a strategy to improve performance with a new sensory skill after some explicit practice. In other words, it is possible that top-down strategies eventually become actively harmful to the process of using a new sensory skill. This would, of course, require more research before any conclusions could be reached.

In conclusion, there can be some similarities to the way that a new sensory skill and native perception function in terms of multisensory interactions. New sensory skills can follow some beneficial patterns like the use of multiple signals to reduce noise or improve speed. This processing can also meet some criteria related to automaticity, suggesting that it can be less like hearing a verbal report about a scene and more like seeing a scene for oneself. There are some notable limitations after this short training: the lack of quantitatively optimal processing, and the lack of forced fusion. The research opens up important questions for future research, including how potential reorganisation of neural sensory processing (Amedi et al., 2017) may support automatic and integrated perception via new signals; how the findings translate to longer training programmes, more complex real-world perceptual-motor tasks; and how they can best be implemented in the design of new devices and approaches for augmenting human perception.

## Supporting information

Raw Data

## Acknowledgments

Thanks to Paula Teo, Rupert Talfourd-Cook, Narjes Kazarooni, Rebecca Leach, and Clara Gallay for help with piloting and data collection. Supported by grant ES/N01846X/1 from the Economic and Social Research Council of the United Kingdom. This project has received funding from the European Research Council (ERC) under the European Union’s Horizon 2020 research and innovation programme (grant agreement No. 820185).

## Full Methods

### Participants

Twelve participants (4 male) took part in this study with an average age of 23.8 (range 20 to 34). The use of 12 participants was taken from the previous study that found cue combination (Negen et al., 2018). An additional two participants were excluded from analysis due to the fact that they failed to learn the audio cue by the second session (female, 20; male, 19). One participant was only able to provide partial data (6 of 10 sessions) due to social distancing measures introduced by the government in March 2020 to counteract the spread of Covid-19. Participants were recruited by word of mouth around Durham, UK. They were paid £10 per session, totalling £100 of compensation. This study was approved by Durham Psychology’s Ethics Board (Reference: PSYCH-2018-12-04). Informed consent was given by participants in writing.

### Apparatus

#### Virtual environment

A custom seascape was created in WorldViz Vizard 5 (Santa Barbara, CA, USA) and presented using an Oculus Rift headset (Consumer Version) (Menlo Park, CA, USA). This seascape contained a large flat blue sea, a ‘pirate ship’ with masts and other items, a virtual chair, and a friendly cartoon whale introduced with the name “Patchy” (see Figure 1). Participants were seated 4.25 m above the sea. The response line (range of possible positions for the whale) stretched out from the bow of the ship and was marked by periodic variations in the colour of the sea. A pair of black bars marked the 10 m and 35 m points, which were the nearest and furthest possible responses. Distances along the line could be judged visually via perspective and height-in plane as well as, in theory, stereo disparity (although stereo information at the distances used is of limited use). Patchy gave written instructions via a white speech bubble (example, Figure 1, far right panel). The sea surface remained still, and the ship did not move. The major advantages of using virtual as compared with real objects was that trials could move much faster, and that we could have control over reliability of visual cues. Full VR via a head-mounted display (rather than a smaller screen) allowed us to immerse people in the virtual environment, avoiding any conflicting spatial information about the real surrounding lab space.

Different trial types also had different kinds of response mechanisms in the virtual world as appropriate. For continuous judgement, there was a grey 3D marker pointed downwards towards the sea (Figure 1). Participants could adjust this along the response line and then enter a response by pressing the A button. For 2-alternative forced-choice (2AFC), 3AFC, and 5AFC trials, the marker would ‘snap’ to the discrete possibilities. For the digit span sections, grey 3D numbers could appear 1m above the sea and 15m out. During the Oddball trials, there were three grey spheres that were 9m away from the ship (Figure 2, top). While the first stimulus played, the left sphere was white; the middle during the second; the right during the third. To respond, participants could use the controller shoulder buttons to select one of the three spheres and then press the A button. For the speed trials, participants would see two small pyramids along the response line (Figure 2, bottom). To indicate that the target was located next to the further pyramid, participants moved the joystick upwards; for the nearer, downwards. As soon as a response was entered by moving the controller joystick, the selected pyramid would grow slightly larger and the non-selected one would turn 50% transparent.

The speed trials also had a special virtual feedback object (Figure 2, bottom). It was a set of cylinders arranged to look like a small stopwatch that floated over Patchy’s head. The maximum time on the clock was 3s. There was a patch on the front that could cover an appropriately sized slice of the clock and change colour. The colour shade varied from red to green based on the time (more red meaning longer).

#### Headset

The Oculus Rift headset has a refresh rate of 90 Hz, a resolution of 1080 × 1200 for each eye, and a diagonal field of view of 110 degrees. Participants were encouraged to sit still and look straight ahead during trials but did not have their head position fixed. The Rift’s tracking camera and internal accelerometer and gyroscope accounted for any head movements in order to render an immersive experience.

#### Audio Equipment

Sound was generated and played using a MATLAB program with a bit depth of 24 and a sampling rate of 96 kHz. A USB sound card (Creative SoundBlaster SB1240; Singapore) was attached to a pair of AKG K271 MkII headphones (Vienna, Austria) with an impedance of 55 ohms.

#### Controller

Participants used an Xbox One controller (Redmond, WA, USA).

### Stimuli

#### Audio

This was a pair of short ‘clicks’ where the time between clicks signals the distance to the target. The audio stimuli were created by first generating a 5ms sine wave 2000 Hz in frequency with an amplitude of 1. The first half-period of the wave was scaled down by a factor of 0.6. An exponential decay mask was created starting after 1.5 periods and ending at 5ms. The exponent was interpolated linearly between 0 and −10 over that period. This was all embedded in 1s of silence, with a 50ms delay before the sound appeared. An exact copy of the sound was added after an appropriate delay, calculating the distance to the target divided by the speed of sound (approximated at 350m/s), then times two (for the emission to go out, and also to come back). With a minimum distance of 10 m, the two sounds (clicks) never overlapped (although it is possible that subjects experienced them as one sound). Real echoes contain more complex information, including reductions in amplitude with distance, but we chose to make delay the only relevant cue so that we could be certain of the information the participants were using. Our stimuli also allowed us to use range of distances at which real echoes are typically very faint, minimizing the scope for participants to have prior experience with them. For the purposes of reaction time, timing began as soon as the 50ms initial delay ended and an actual sound began playing.

#### Visual

This stimulus is essentially a type of coherent motion or motion integration stimulus. There were 100 black blocks with a side length of 0.05m. From the participant’s perspective, the x axis is left/right and the z axis is near/out. At the beginning of the stimulus, the boxes were spread evenly from 10m to 35m along the z axis. They were placed uniformly randomly from −0.5m to 0.5m along the x axis. Over the course of 250ms, beginning as soon as they appeared, each box moved 20% of the way towards the target. As soon as the 250ms ended, the blocks disappeared. Any inaccuracy represents internal noise, since perfectly estimating the block trajectories’ convergence point (or even perfectly extrapolating the remaining 80% of any one block’s trajectory) would perfectly localise the target. There is also a longer version of this stimulus, intended to make it more reliable, for certain trial types. This is the same, except that blocks move 40% of the way towards the target over the course of 500ms. For the purposes of reaction time, timing begins as soon as the blocks appear.

#### Digit Span

Six digits were selected randomly. A text-to-speech voice read out the six digits in order at a rate of one per second.

### Procedure

There were 10 sessions, each with up to 300 trials. Each session was done on a different day and allowed to span up to 10 weeks. Breaks were given whenever requested. The goal and types of trials were slightly different for almost every session. Figure 8 gives an overview of which trial types each session used and in what order. The following text explains each session in detail. Whenever a session uses a new trial type that has not yet been used, that trial type will be explained under a subheading.

**Figure 8:**
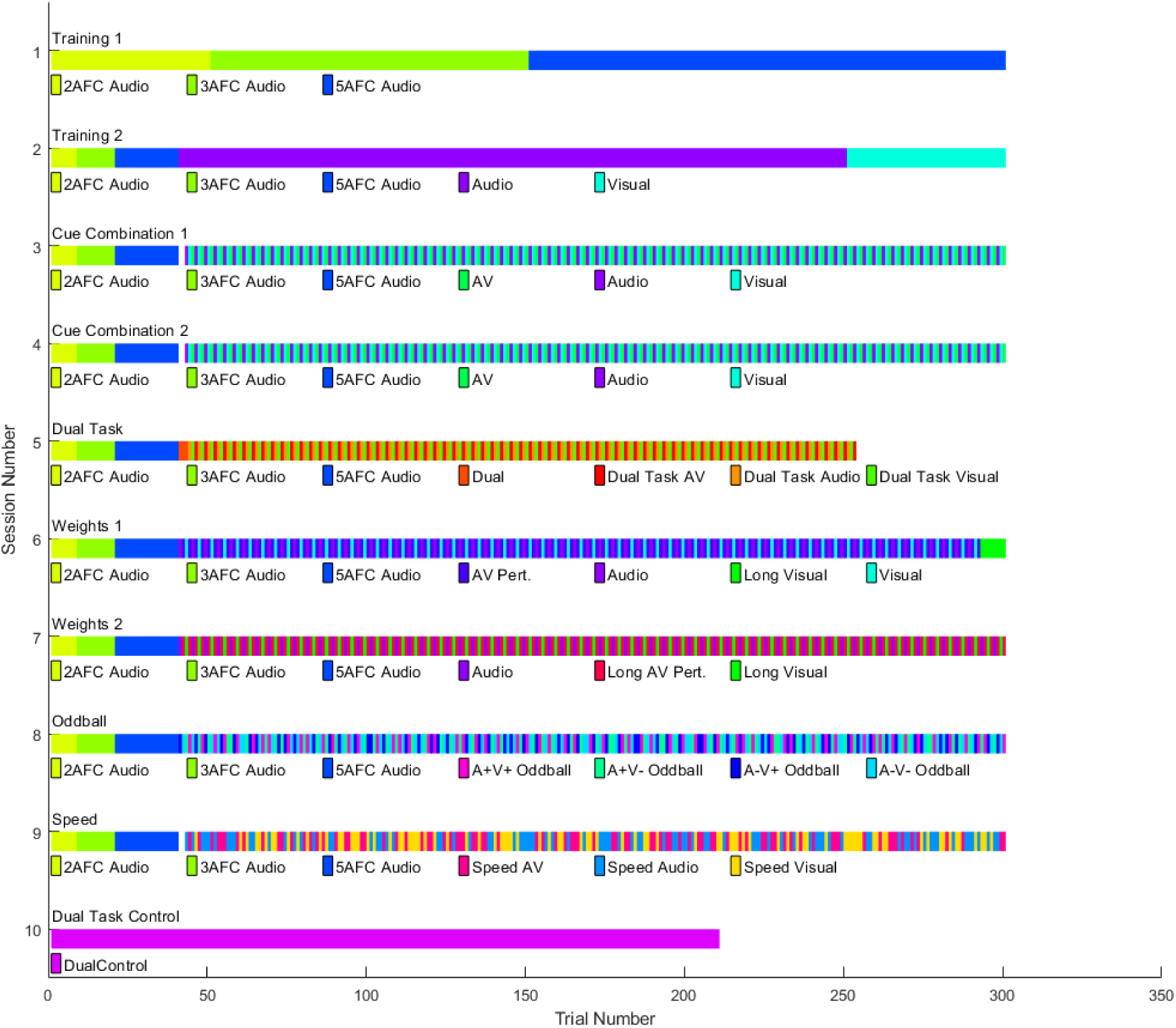
Reference guide for the different sessions and trial types. See *Procedure* for details of each trial type and an explanation of each session’s purpose.

#### Training 1 (Session 1)

The aim of the first session was to train the participants to use the audio cue. No data from this session were analysed. The session consisted of 50 trials of 2AFC Audio, followed by 100 trials of 3AFC Audio, and then 150 trials of 5AFC Audio. Targets were used as evenly as possible and in a random order.

##### 2AFC Audio

2AFC stands for two-alternative forced choice. With an audio stimulus, participants judge if the target was 10m or 35m away. Before the first 2AFC Audio trial, Patchy would demonstrate the difference by alternatively appearing 10m and 35m away while the matching audio stimulus played. This demonstration cycled six times. During testing trials, feedback was given: if the correct choice was made, Patchy would appear and make a ‘nodding’ motion. If the incorrect choice was made, Patchy would appear at the correct location and move his head left and right in a ‘no’ gesture.

##### 3AFC Audio

With an audio stimulus, participants were required to judge if the target was 10m, 22.5m, or 35m away. Before the first 3AFC Audio trial, the three distances were demonstrated in four cycles. Feedback was the same as 2AFC Audio.

##### 5AFC Audio

With an audio stimulus, participants were required to judge if the target was 10m, 16.25, 22.5m, 28.75, or 35m away. Before the first 5AFC Audio trial, the five distances were demonstrated in three cycles. Feedback was the same as 2AFC Audio.

#### Training 2 (Session 2)

Session 2 was further training with the audio cue, a test of the Cue Learning (Preliminary) Hypothesis, and an introduction to the visual stimulus. First the participant was presented with 8 trials of 2AFC Audio, followed by 12 trials of 3AFC Audio, and then 20 trials of 5AFC Audio which served as a reminder of the previous training. Targets were used evenly in a random order. For the following text, that set of 40 trials will be referred to as ‘the warmup block’ for brevity. The warmup block is not analysed here or in any session. After the warmup block, there were 210 Audio trials. These were used to test the Cue Learning (Preliminary) Hypothesis, which predicts a significant correlation between target and response, and to give further training with the audio cue. Next there were 50 Visual trials, again spread evenly on a log scale and presented in a random order. This was to ensure that the participants were adequately familiarised with the visual cue and its use before the tests of cue combination (i.e. we would not want people to fail to combine the audio and visual cue due to unfamiliarity with the visual cue). These Visual trials are not analysed.

##### Audio

Participants were only given an audio cue (the new sensory skill) to use. They were required to judge the distance to a target along a continuous line stretching from 10m to 35m. They used a joystick on the controller to move a marker to their estimated target location and pressed the A button. Patchy would appear and give them feedback in terms of percentage. For example, a target of 20m and a response of 18m would get the feedback “-10.0%”.

##### Visual

During visual trials, only the visual stimulus is presented.

#### Cue Combination 1 (Session 3)

Sessions 3 to 5 were designed to test the Precision Hypothesis (Cue Combination). This hypothesis states that variable error will be lower in the AV trials than with the best single cue (i.e. whichever of Audio or Visual has the lower variable error). These sessions also provided a basis for estimation of the optimal possible precision for the Optimal Weight (Cue Combination) Hypothesis. After the warmup block (not analysed), there were 86 Audio, 86 Visual, and 86 AV trials. They were presented in the order of one Audio trial, then one Visual, then one AV, then one Audio, and so on. The targets were spread evenly on a log scale and random in order.

##### AV

During AV trials, both the audio and visual stimuli are presented. For this trial type, the two always agree perfectly (i.e. there is no offset in the distances they signal) and begin at the same time.

#### Cue Combination 2 (Session 4)

We repeated the procedure from session 3 in session 4 in order to gather more data and provide greater statistical power for the Precision Hypothesis (Cue Combination) and the Optimal Weight (Cue Combination) hypothesis.

#### Dual Task (Session 5)

Session 5 served several purposes at once. The primary purpose was to test the Parallel to Verbal (Automaticity) hypothesis. This states in part that performance on the Audio, Visual, and AV trials is independent of verbal working memory and thus will not be impaired by a simultaneous verbal working memory task. This set of trials also provided further data for the Precision Hypothesis (Cue combination) and the Optimal Weight (Cue Combination) hypothesis. After the warmup block (not analysed), there were three Dual Task Practice trials (not analysed). Following this, there were 70 Dual Task Audio, 70 Dual Task Visual, and 70 Dual Task AV trials. These were presented in the order of one Dual Task Audio, one Dual Task Visual, one Dual Task AV, one Dual Task Audio, and so on. The targets were spread evenly on a log scale and random in order.

##### Dual Task Practice

Participants were presented audio visually with six random digits. There was then a delay of 2.0 seconds before they completed a memory probe. For the probe, they were presented with a visual display containing five of the six digits. The missing digit is replaced with a question mark. To identify the missing digit from memory, they used the controller to cycle through digits 0-9 and clicked to indicate their response. No feedback was given.

##### Dual Task Audio

The participants were presented with the random digits before hearing an audio stimulus (i.e. digits then clicks). They were then required to judge the distance to a target (continuously 10m to 35m), and then complete a memory probe. No feedback was given on the memory probe. Feedback on the distance judgement was given after completing the memory probe.

##### Dual Task Visual

This task was a replication of Dual Task Audio, but the auditory stimulus was replaced with the visual.

##### Dual Task AV

This task was similar to the Dual Task Audio, but with the addition of a visual stimulus for judging distance.

#### Weights 1 (Session 6)

Sessions 6 and 7 were designed to test the Optimal Weight (Cue Combination) hypothesis. This required us to estimate the actual weights placed by the participant on each cue. It also required that we estimate the optimal weight each cue should be given so that optimal weights can be compared with the actual weights. After the warmup block (not analysed), there were 63 Audio trials, 63 Visual trials, and 126 AV Perturbation trials. The AV Perturbation trials allowed us to estimate the weight given to each of the two cues via multiple regression. The Audio and Visual trials were performed in order to estimate the precision (1/variance) of responses with each single cue, which then determined the optimal weight to give each one (higher precision means more weight). These were presented in the order of one Audio trial, one AV Perturbation trial, one Visual trial, one AV perturbation trial, one Audio trial, and so on. There were 63 targets spread evenly on a log scale. Each trial was used once for the Audio trials, once for the Visual trials, and twice for the visual stimulus of the AV Perturbation trials. For the AV Perturbation trials, the audio distance was +/- 10% of the visual distance, with the sign chosen randomly unless one choice would place the audio stimulus outside of the response range of 10m to 35m. Targets were presented in random order. At the end of the session, 8 Long Visual trials (not analysed) were presented to introduce participants to the new visual reliability for the next session.

##### AV Perturbation

The participants were given an audio cue and a visual cue that differed from each other by 10%. To be very specific, once a location was selected, the visual cue indicated the true location while the audio cue signalled a distance that was plus or minus ln(1.1)=0.0953 on a natural log scale. The sign was chosen randomly unless one sign would place the audio stimulus outside the response range (10m to 35m). The participant made a judgement of the distance to the target along a continuous line. No feedback was given.

#### Weights 2 (Session 7)

Session 7 was a further test of the Optimal Weight (Cue Combination) Hypothesis. For this session, the longer version of the visual stimulus was used. We expected this to lead to higher precision in visual judgments, and therefore to change the optimal-predicted weighting towards vision. After the warmup block, there were 65 Audio trials, 65 Long Visual trials, and 130 Long AV Perturbation trials. The scheme for trial order and target placement mirrored Session 6. This again allowed us to estimate the actual weight and optimal weight given to each.

##### Long Visual

Participants were presented with a visual stimulus lasting twice as long as the usual Visual trials (500 vs 250ms) and were required to estimate the location on a continuous line ranging from 10m to 35m.

##### Long AV Perturbation

This was a replication of the AV Perturbation condition, but with the long visual stimulus previously described in place of the usual short visual presentation. There was no feedback given.

#### Oddball (Session 8)

Session 8 was designed to test the Forced Fusion (Automaticity) hypothesis. The standard method for testing this is an oddball task with congruent versus incongruent trials, with forced fusion inferred if the incongruent trials are more difficult (Hillis et al., 2002). After the warmup block (not analysed), there were 65 A+V+ Oddball trials (congruent), 65 A-V-Oddball trials (congruent), 65 A-V+ Oddball trials (incongruent), and 65 A+V-Oddball trials (incongruent). These were presented in a random order. The standards were spread evenly from 15.7m to 22.3m. Targets were selected through a staircase method, described below.

##### A+V+ Oddball

A standard distance was chosen along the line from 15.7m to 22.3m. An oddball distance was generated that differed from the standard by a certain amount. The participant was presented with three sets of AV stimuli. The standard was played twice and the oddball once. Each stimulus lasted 1.0s. The task was to select the oddball, which was randomly chosen to be one of stimuli 1-3. In the instructions, this was specifically phrased as: “Two are the same. One is different. Pick the odd one out.” In an A+V+ Oddball trial, the audio and visual components of the oddball presentation both signalled a further distance than the standard. The difference between the standard and oddball was controlled by a staircase procedure, with each oddball trial type having a separate staircase. The difference was on a log scale. Each staircase began at 0.2. With every correct response this was multiplied by 0.9. With every incorrect response this was multiplied by 1/(0.9^3). This was capped at 0.4. For example, suppose a 20m standard and a difference of 0.2 on a log scale. The oddball would signal a distance of e^(ln(20)+0.2) = 24.4m. No feedback was given.

##### A-V-Oddball

These trials were the same as the A+V+ Oddball trials, with the exception that the audio and visual stimulus in the oddball presentation signalled a distance that was nearer than the standard. No feedback is given.

##### A+V-Oddball

This was like an A+V+ Oddball, with the exception that the audio stimulus in the oddball presentations signalled a distance which was further than the standard and the oddball’s visual stimulus signalled a distance that was nearer than the standard.

##### A-V+ Oddball

This is like an A+V+ Oddball, with the exception that the audio stimulus in the oddball presentations signalled a distance nearer than the standard and the visual stimulus in the oddball presentations signalled a distance that was further than the standard.

#### Speed (Session 9)

Session 9 was designed to test the Redundant Signals (Automaticity) Hypothesis. This hypothesis suggests that the new audio cue will facilitate faster responses alongside the visual cue than could be achieved with the visual cue alone. After the warmup block (not analysed), there were 83 Speed Audio, 83 Speed Visual, and 83 Speed AV trials. These were presented in random order. Targets were spread evenly on a log scale. Incorrect alternatives differed by 40%. Since the decisions were relatively easy, the instructions instead stressed the importance of speed for these trials: “This one is about SPEED!” and “Ready? Remember, FAST!”.

##### Speed Audio/Visual/AV

The participant was shown two possible distances to choose which differed by 40%. They were marked by small pyramids appearing. After the pyramids appeared, but before the stimulus began, there was a delay with a randomized length: 0.5 seconds plus an amount drawn from exponential distribution with a rate of 3. This is capped at 2.5 seconds. The stimulus (Audio, Visual, or AV) indicated one of the two possible distances. The participant was required to move the joystick upwards to indicate the further distance or downwards to indicate the nearer distance. Feedback was then given in the form of a small clock displaying their time if they were correct, otherwise, Patchy appeared and slowly shook his head in a ‘no’ gesture. Several randomly selected trials were also designated as ‘catch’ trials where the delay was increased by 2.5 seconds but the trial was otherwise the same.

#### Dual Task Control (Session 10)

Session 10 (along with session 5) was designed to test the Parallel to Verbal (Automaticity) hypothesis. These trials were designed to measure performance on the verbal working memory task without the need to make a distance judgment. Participants still used the joystick to place the marker but could now see the location of the target (Patchy) directly rather than having to infer it via uncertain audio or visual cues. There was no warmup block. There were 210 Dual Task Control trials.

##### Dual Task Control

Participants heard six random single digit numbers. Following a delay of 2.0 seconds, they saw Patchy appear. They then used the joystick to place the marker onto Patchy and press A. They then completed a memory probe (i.e. they were presented with the previously heard six digit sequence visually, with a question mark in place of the probe digit). They then use the controller to select the correct number to place into the sequence. No feedback is given.

### Data Processing and Analysis Plan

The procedure detailed above resulted in a total of 2,857 trials for each participant. In this section, we describe how we extracted measures from this dataset and analysed them.

#### Cue Learning (Preliminary) Hypothesis

For the audio trials from session 2, for each participant the target and response were transformed onto a logarithmic scale. This was done to account for Weber’s law (Getty, 1975). The data were then analysed for a correlation between the targets and responses. A positive correlation in an individual participant was interpreted as evidence for that participant learning how the audio cue works. For comparison to expert echolocators, we also estimated a weber fraction for each participant. This was be done by creating synthetic two-alternative forced-choice (2AFC) trials from the audio trials. We assumed that if an estimate when given one target is further than an estimate given another target, then the participant would have chosen the first target as further in a 2AFC task. This method is not as ideal as actual 2AFC data but should at least give some indication of how far the overall performance is from expert performance.

#### Parallel to Verbal (Automaticity) Hypothesis

For this and further analyses, we needed to be able to calculate the variable error (i.e. to estimate the amount of perceptual noise in the judgements, separated from systematic distortions). This was done with the data from sessions 3-5, using the trial types that involve a continuous judgement of distance (Audio, Visual, AV, and their Dual Task variants). First, we trimmed outliers. This was done by calculating the standard deviation of responses minus targets, on a logarithmic scale, for all the data being used here. This was 0.23. Any trial with an error greater than three times this (0.69) was excluded as an outlier. This corresponded roughly to any response that was more than twice the target distance or less than half the target distance. This excluded 145 trials (1.66%). Second, we trimmed the outer 10% of targets to remove any heavy distortion due to the bounds on the response range. Unfortunately, even after this, there was still a central tendency bias; the slope of targets regressed onto targets was less than 1.0 on average, t(107) = −4.07, p < 0.001, d = −0.39. We therefore had to employ a somewhat unusual method of calculating the variable errors to avoid biasing the results.

The next steps were done separately for each participant, session, and trial type. Third, we regressed the responses onto the targets, with both expressed on a logarithmic scale. This resulted in a slope of the regression line and a set of residuals (errors from the regression line). Fourth, the variable error was calculated by finding the standard deviation of the residuals, then dividing that by the slope, with the slope capped at 1.0. This was done because a central tendency bias could mask the amount of perceptual noise. For example, suppose a participant takes their actual perceptual estimates and moves them 50% of the way towards the centre of the response range. This would reduce the standard deviation of the residuals by 50% without changing the perceptual noise at all. It would also result in a slope of 0.5. Dividing the standard deviation of the residuals by the slope recovers the original perceptual noise. This measure is an attempt to directly capture the kind of perceptual noise that distraction should increase and cue combination should reduce, separate from systematic distortions like central tendency bias. This results in a 12 (participants) x 3 (session 3, 4, or 5) x 3 (Audio, Visual, or AV) matrix of variable errors. These variable errors are taken as the estimate of perceptual noise.

Since this is an unusual variable error calculation, we will comment further on its properties to aid understanding. It is not just measuring raw error; it parses away the systematic (linear) error and measures the remaining random error. As it happens, our variable error measure here is mathematically and conceptually related to the Pearson’s R statistic. You could divide all the responses by 10 (or any other number >1) and it would not change our variable error measure. You could add 10 (or any other number) to all the responses and it would not change our variable error measure. We assume that such systematic (linear) distortions are not due to perceptual noise, but instead are better explained by things like a central tendency bias. In the special case where the correlation between targets and responses is exactly 1.0, our variable error measure is guaranteed to be zero. As it happens, the results for this hypothesis remain the same if the correlation between target and response is substituted for our variable error measure in the rest of the analysis (except in the opposite direction, with a lower variable error corresponding to a higher correlation). The same is true for the Precision Hypothesis (Cue Combination) presented later. We are using this variable error measure instead of a correlation mainly because it enables the calculation of optimal weights for the Optimal Weight Hypothesis. In the special case where there are no linear distortions, our measure is the same as the standard deviation of the raw log errors (response minus target on a log scale). While this measure has not been used by us before, we foresee using it as our standard way of calculating variable error in our future projects with continuous responses.

The rest of the process has two sections. First, we wanted to see if the digit span task affected distance judgements. The variable error in sessions 3 and 4, without the digit span task, was compared against the variable error for session 5, with the digit span task. This was done as a 2 (session type) x 3 (Audio, Visual, or AV) repeated-measures ANOVA. To facilitate this, the variable errors for sessions 3 and 4 were averaged together over sessions, within participants and within trial types. Second, we wanted to see if the distance judgement affected accuracy on the digit span task. To do this, for each participant we computed the average of the percent of correct memory probes in Dual Task Audio, Dual Task Visual, and Dual Task AV. This was a 12-participant vector of percent correct scores. This was compared with a paired t-test against the percent of correct memory probes in session 10 (i.e. Dual Task Control).

#### Redundant Signals (Automaticity) Hypothesis

This section involves mean reaction times and accuracy rates. Since reaction times often have many outliers, we used the 90% trimmed mean for all responses. For each participant, we calculated the 90% trimmed mean of reaction times for Speed Audio trials, for Speed Visual trials, and for Speed AV trials. This resulted in an 11 (participant) x 3 (Speed Audio, Speed Visual, or Speed AV) matrix of average reaction times. We also calculated the percent correct for Speed Audio trials, Speed Visual trials, and Speed AV trials.

First, we calculated the mean reaction time for Speed Audio trials along with a 95% confidence interval. Second, we compared the mean percent correct in Speed Audio trials against the chance rate (50%) with a one-sample t-test. Third, we wanted to compare the average reaction time with each cue alone (i.e. Speed Audio and Speed Visual) versus the average reaction time in Speed AV trials. We compared Speed Audio versus Speed AV with a paired t-test and then compared Speed Visual versus Speed AV with another paired t-test. Fourth, we wanted to check that any speed gains could not be explained purely by adoption of a different speed-accuracy trade-off, which would be evident in a loss of accuracy. Two paired t-tests were again used to compare accuracy in Speed AV against Speed Audio and then Speed Visual.

#### Forced Fusion (Automaticity) Hypothesis

There is unfortunately no standard model for fitting and analysing oddball data. It is not a 2AFC task and there is no consensus theoretical reason to believe that performance will follow a specific curve. We therefore analysed these data in two ways. The first remains very close to the raw data and does no fitting. For this, we used the average congruent deviation and the average incongruent deviation. On each oddball trial, there are two standard choices and the oddball choice, which deviates from the two standard choices. Since these trials used a staircase design, the average deviation is an index of performance. The deviations were expressed on a log scale (i.e. on the same scale as the staircase). The average congruent deviation was the average deviation in A+V+ trials and A-V-trials. The average incongruent deviation was the average deviation in A+V-trials and A-V+ trials. A paired t-test was used to compare congruent and incongruent average deviations. Forced fusion predicts greater deviations for incongruent stimuli. In addition, if this measure is capturing the ability to discriminate the oddball from the standard in a reasonable fashion, then we expect it to capture individual differences. To check that this is true, we examined the correlation between the congruent versus incongruent outcome measures.

For the second method, we adapted a typical 2AFC model based on the Gaussian cumulative density function (CDF). We assumed that the probability of a correct choice is:

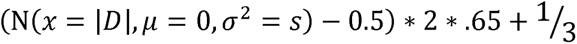

Where N() is the normal CDF, *D* is the deviation of the oddball from the standard on a log scale, and *s* is the fitted parameter. This equation has a minimum of 1/3 where D=0, has a limit of 0.98333… as *D* moves away from 0, and is monotonically increasing as *D* moves away from zero. While there is little in the way of deep theory behind this equation, it at least satisfies some basic properties that we would like: the probability of a correct response is at least equal to chance guessing (1/3), at most near 100% but with a small overhead for lapses, and increases as the oddball becomes increasingly different from the standard. With smaller values of *s*, the equation moves more steeply towards its upper limit as a function of *D*. We fitted the value of *s* to each participant by maximizing the likelihood of the observed data. We then solved for the value of *D* that results in a probability of 2/3 for a correct response and took that as the fitted threshold.

#### Precision (Cue Combination) Hypothesis

Here, we wanted to compare the variable error for the best single cue against the variable error for AV trials. The variable error for the best single cue was taken as the variable error for either audio or visual, whichever was lower. This could be different for each combination of participant and session. This resulted in a 12 participants x 3 sessions x 2 (best single, AV) matrix. The data were then analysed with a repeated-measures ANOVA (best single versus AV). The section above regarding the Parallel to Verbal (Automaticity) Hypothesis describes the calculation of variable errors.

#### Optimal Weight (Cue Combination) Hypothesis

For this we needed the optimal variable error (i.e. perceptual noise as a standard deviation) for each participant during sessions 3-5, the weights given to each cue during sessions 6 and 7, and the optimal weights to give each cue during sessions 6 and 7. The optimal variable error for sessions 3-5 was found with the formula:

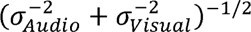

The variable errors are standard deviations. This formula transforms them to precisions (1/variance), adds them, and then transforms them back to standard deviations. This results in a 12 (participant) x 3 (session 3, 4, or 5) matrix of optimal variable errors. This was compared to the AV variable error for each participant in sessions 3-5 with a repeated-measures ANOVA.

The optimal weight to give each cue was calculated separately for each participant and session (6 or 7). The optimal weight to give the visual cue (Ernst et al., 2016) is

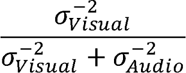

The optimal weight to give the audio cue is one minus the optimal visual weight. Standard deviations were found for the audio trials and the visual trials in the same manner as described above (Parallel to Verbal hypothesis). This resulted in an 11 (participant) x 2 (session 6 or 7) x 2 (Audio or Visual) matrix of optimal weights. (The twelfth participant only provided data for sessions 1-5.) These are compared against the observed weights, described below.

The observed weights were found by multiple regression. This was done separately for each participant and session (6 or 7). The natural logarithm of the responses was the outcome variable. The distance signalled by the audio and visual cue, both on a logarithmic scale, were the predictors. The weights were taken to be the estimated beta value for each predictor. In four cases where this was below zero or above one, they were adjusted to be zero or one. Note that this procedure does not always result in observed weights that sum to exactly one, but did tend to remain close to this (average of 0.95). This results in an 11 (participant) x 2 (session 6 or 7) x 2 (audio or visual) matrix of observed weights.

Observed visual weights were compared to optimal visual weights with a repeated-measures ANOVA. The same was done for audio weights. The correlation between observed visual weights and optimal visual weights was also analysed; again, the same for audio. Finally, a paired t-test was used to see if the visual weight was higher in session 7, which had a longer (i.e. more reliable) visual stimulus, than in session 6, which had the normal (i.e. less reliable) visual stimulus.

## Appendix – Cross Validation for Cue Combination

Results indicate that participants combined the native visual cue and the new audio cue to judge distance. However, results also point towards sub-optimal combination – specifically a tendency to over-weight the visual cue and under-weight the audio cue. This appendix supplements those analyses with a cross-validation approach to investigate the possibility of convergent evidence for this interpretation. In summary, the results presented here agree with the main text.

### Data Used

For these models, we use the trials that involve a distance judgement, starting with session 3 (after training). This includes the dual task trials, the perturbation trials, and the trials with a longer visual cue. The 2AFC trials, 3AFC trials, 5AFC trials, speeded task trials, control task trials, and oddball task trials are not used. In session 6, where there are 8 visual-only trials with the long visual cue and 83 trials with the shorter visual cue, only the shorter ones are used. The data from the final participant with a partial dataset are used where possible. Within each session and participant, the nearest 5% of targets and the furthest 5% of targets are excluded. Each session is modelled separately i.e. the same participant can have completely independent parameters in sessions 3, 4, 5, 6, and 7.

### Additional Background

For readers who are unfamiliar with cross-validation and Bayesian cue combination, here we provide further background to the below formulae.

#### Bayesian Weights

In the Bayesian framework, whenever there are two sources of information that both contribute to the same decision, each estimate is given a weight for a weighted average. Suppose we are trying to judge the distance to a target that is 10m away. The visual system might estimate that it is 11m and the audio system might estimate that it is 8m. Suppose that an audio cue has a perceptual variance of 3m (i.e. over the course of many estimates, the error in the perception of the cue has a variance of 3m). That means it has a perceptual precision of 1/3 = 0.33. Suppose that a visual cue has a perceptual variance of 2m. That means it has a perceptual precision of 1/2 = 0.5. We can find the weight by calculating the proportion of summed perceptual precisions. For our audio cue, that would mean 0.33/(0.5+0.33) = 40%. For the visual cue, that would mean 0.5/(0.5+0.33) = 60%. For our distance judgement, we make our final estimate by multiplying 60% (visual weight) times 11m (visual estimate) plus 40% (audio weight) times 8m (audio estimate). That is 0.6*11+0.4*8 = 9.8m, which is more accurate than either estimate alone. This pattern will hold more often than not: the weighted average, with this method of making weights using these precisions, is more accurate than either estimate alone. In this situation, we could write the weight for the audio cue like so:

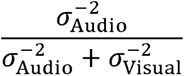

In the formulas under *Models*, terms like this, with σ on top and bottom of a fraction, are weights based on precisions. Since audio is on the top, we can see that this is the weight for an audio cue.

Bayesian weights also apply beyond sensory cues. Prior distributions can get the same kind of weight with the same kinds of mechanics. A prior distribution, formally, is the belief that some locations are more likely to contain targets than others. If the prior distribution is Gaussian, then Bayesian mechanics will simply give the mean of the prior a weight and make it part of the weighted average. Entering a prior distribution pulls responses towards the prior mean. This can be used, for example, to model a bias towards the centre of the response line regardless of why exactly it happens. The only important mathematical difference between a sensory cue and a prior mean is that the prior mean doesn’t vary from trial to trial. This is because the prior mean has nothing to do with the target and has no sensory noise.

We can also weight more than two things this way. For example, suppose we want the weight for an audio cue when there is an audio cue, a visual cue, and a prior distribution. It is:

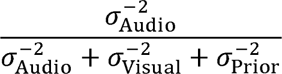

It is just a matter of placing everything we are averaging at the bottom of the fraction. In principle, this could be extended to four, five, and so on (though we will only need three here).

Note that this section is only strictly true if we are working with normal distributions. Here, we only work with normal distributions.

#### Lapse Mechanics

Cross-validation can give extremely strange results if there are outliers in the data and the models under consideration do not have any way of addressing them. In extreme cases, the final score of a model can be almost entirely determined by its ability to explain a single outlier data point. It is therefore important to build in a kind of ‘safety valve’ where no single data point can have an extremely low probability. The best way to do this will depend the model and the nature of the data.

Here we do this by building in a lapse rate of 2%. Basically, we ask what a participant would do if they were responding at random on a certain trial. In this case, their response would follow a uniform distribution across the entire response line. The nearest possible response is 10m. The furthest is 35m. We are working with the natural logarithm of targets and responses, so the full range is ln(35)-ln(10). To spread 2% of the probability over this range, we can add this term to the probability of every response:

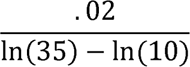

This results in a value of approximately 0.016. Whenever you see this term in the models below (with .02 on top and natural logarithms below), it is the lapse mechanic. In other words, the models get 98% of the probability to use for fitting, but 2% is always reserved and spread evenly over the response range as a lapse mechanic. No single data point can have a probability below about 0.016, so we will avoid a situation where any single data point has an extremely low probability and dominates the final score by itself.

#### Standard Deviation of a Weighted Average

Suppose we draw two numbers, x and y, from two distributions, X and Y. Suppose their standard deviations are σ_*X*_ and σ_*Y*_. Suppose we then take a weighted average of x and y, giving them the weights W_X_ and W_Y_. If we do this over and over, the standard deviation of the weighted average will be

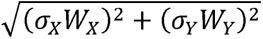

Below, we can see this being used in Equations 4a and 7a (with the audio cue as X and the visual cue as Y).

### Models

The three models are Single Cue, Optimal, and Audio Confidence. Common to all three, we use the natural logarithm of the cues and the responses. Also common to all three, there is a lapse mechanic and a mechanic for the application of a prior. The lapse reserves 2% of the probability to be spread evenly along the response line. The prior allows for the participant to have their responses biased systematically towards a particular point on the line. While this is modelled explicitly as a prior, the mathematics can also mimic a more general bias e.g. a central tendency bias. Non-lapse trials are modelled as a normal distribution.

#### Single Cue

Conceptually, single cue means that participants only use a single cue when two cues are available. The analysis in the main text suggests that this model is not favoured, at least for most participants. This is an important alternative because it represents the way that young children, who are still learning to use their native senses for the first time, deal with similar multisensory situations (Burr & Gori, 2012). It is also the way that participants were previously shown to were shown to address the simultaneous presentation of a trained new sensory cue and a native vestibular cue (Goeke et al., 2016).

Mechanically, this model has five parameters:

1. σ_Audio_ the standard deviation of perception around audio cues. This (and the next two) are constrained to be positive values only.
2. σ_Visual_ the standard deviation of perception around visual cues.
3. σ_Prior_ the standard deviation of a prior that the participant applies.
4. μ_Prior_ the mean of the prior that the participant applies. This value is not constrained.
5. *V* the probability that the participant uses the visual cue. 1-V is the probability that they use the audio cue. This is constrained to be at least zero and at most one.

The modelled distribution of the single-cue trial types follows:

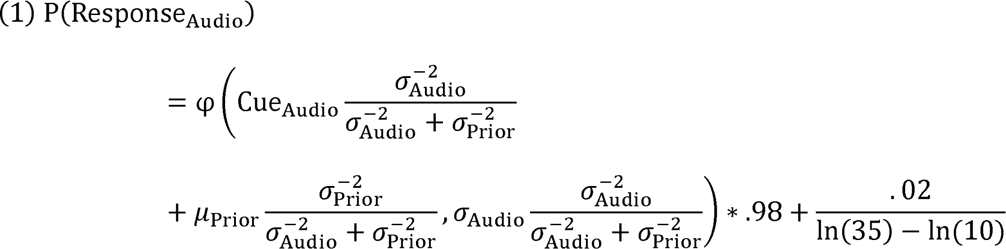

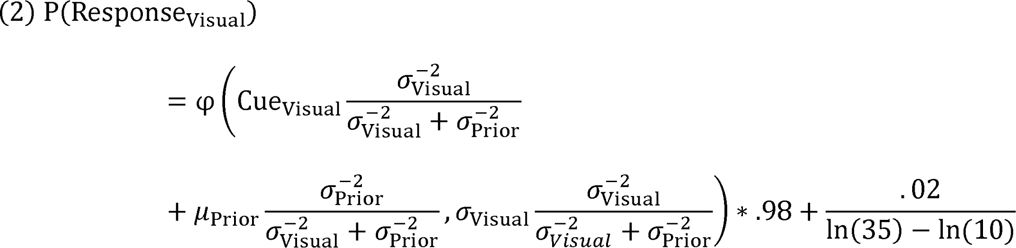

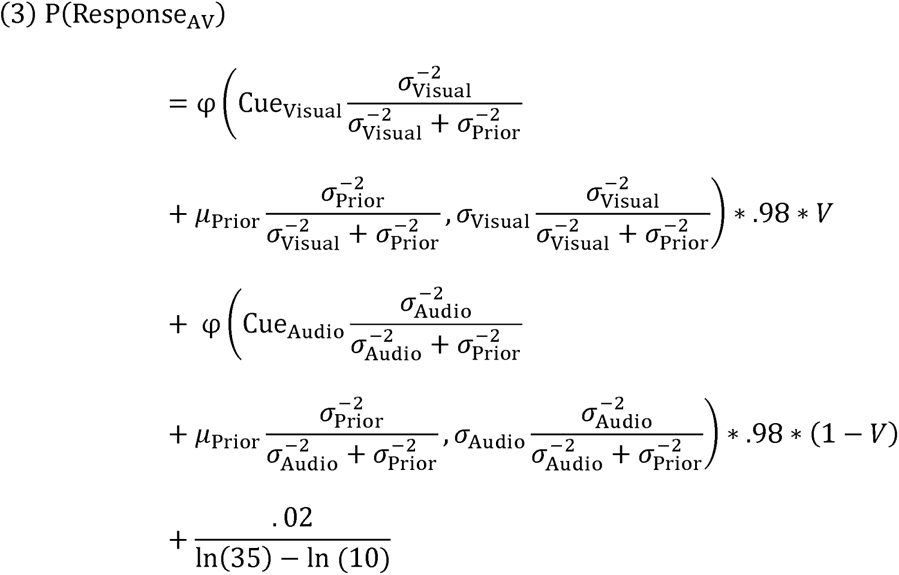

where φ is the probability density function of the normal distribution, parameterized with a mean then a standard deviation, and ln() is the natural logarithm. In the first two equations, we can see that the φ() term is multiplied by 0.98 and that a term spreading 2% of the probability across the response range (from 10m to 35m) is added. That is the lapse mechanic. We can also see that the mean response is a weighted average of the cue and the µ_Prior_ term. The standard deviation is also adjusted for the possibility that the cue has a weight below one. Those are the mechanics of the prior that the participant applies. In equation 3, we see that the first φ(), term depends on the visual cue and is multiplied by *V*, while the second φ() term depends on the audio cue and is multiplied by (1-V). When V is set to 1, only the visual cue is used. When V is set to 0, only the audio cue is used. When V is in intermediate, the participant switches back and forth between the two available cues at random, but does not use both on the same trial.

#### Optimal

The optimal model means that participants use the two cues together optimally. The analysis in the main text again suggests that this model is not favoured for most participants. This is an important alternative because it reflects the way participants deal with many multisensory tasks that only involve native cues (Pouget et al., 2013).

Mechanically, this model uses the same parameters as Single Cue, except without the final *V* parameter. It also uses the same equations for the audio-only and visual-only trials (i.e. equations 1 and 2). It differs in the equation for the AV trials:

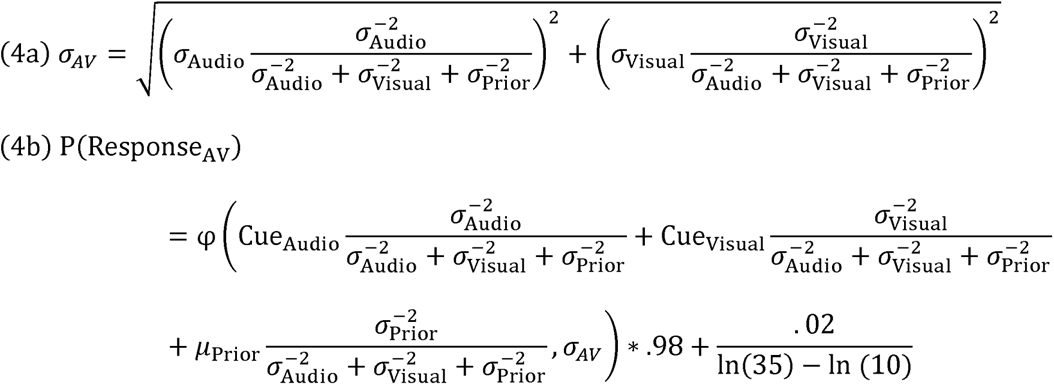

Here we see that the mean response is a weighted average of the audio cue, the visual cue, and the prior mean. The standard deviation is a pool of the audio standard deviation times the audio weight as well as the visual standard deviation times the visual weight. These weights are optimal.

#### Audio Confidence

Conceptually, Audio Confidence allows a participant to under-estimate the reliability of the audio cue. Otherwise, it is equivalent to the optimal model. The analysis in the main text favours this model. If this model is not favoured here, at least for most participants, that would cause us to carefully re-examine the interpretation given in the main text.

Mechanically, this model uses the same first four parameters. It also uses an additional free parameter *C* that is a factor to reflect under-estimation of audio precision. *C* is constrained to be at least zero and at most one. *C* can be inferred by looking at the AV trials, especially the ones with a perturbation, to see if they over-rely on the visual cue. Equation 2 is still used for the visual-only trials. The other two equations are

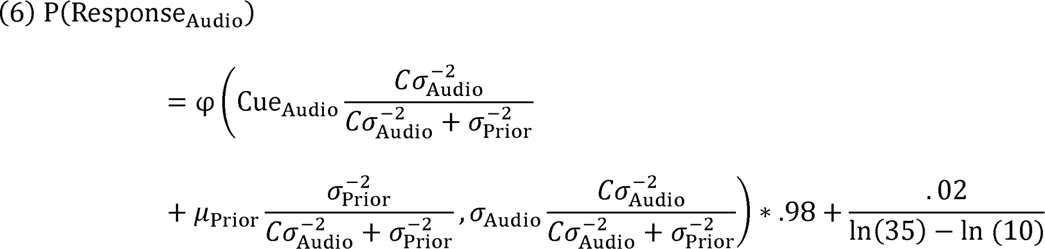

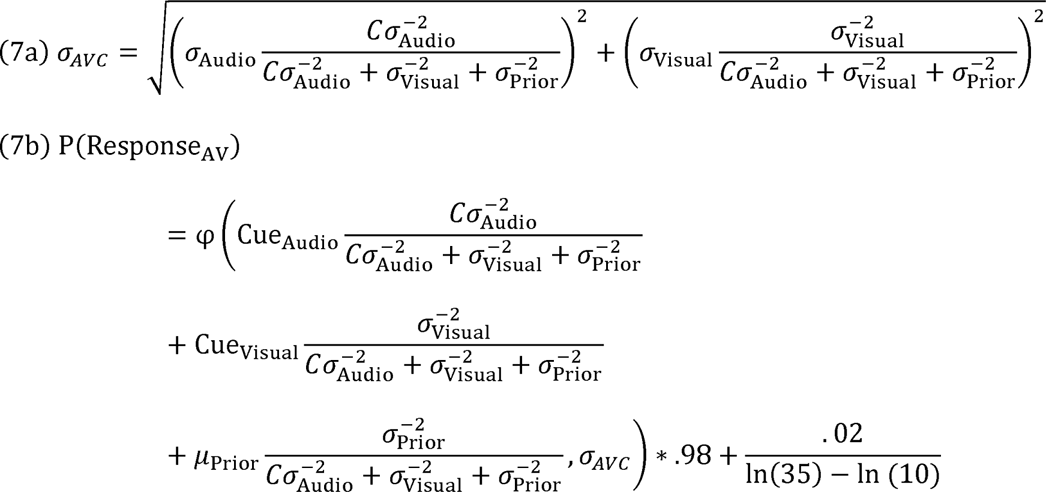

Basically, the actual precision of the audio cue, 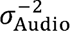, is multiplied by *C* in every instance where we are calculating a weight.

One could equivalently state that the Optimal model is the Audio Confidence model with C constrained to equal one.

### Cross Validation Methodology

Cross validation is a method that tries to favour models that fit well but also guards against overfitting. It is not just a measure of fit; one model can have a higher maximum likelihood estimate fit while having a lower cross-validation score. Conceptually, it is like we pretend that some of the data have already been observed and that some of the data are going to be collected later. The procedure tests how well the model can use the data we already have to predict the future data. There is no p-value or Bayes factor – rather, a very general measure of how well the model functions as a tool for making future predictions.

The procedure is done separately for each participant and each session. The data are cut into *k* slices (i.e. smaller sets of data). Here we used *k*=10. On each of *k* rounds, one slice is designated as the testing data. The other *k*-1 slices are designated as the training data. The training data are fit in the sense of finding the parameter values that maximize the likelihood of the training data. We then calculate what joint probability those parameters assign to the testing data. The joint probability of that slice is stored. This is done once for each slice (i.e. each slice is used once as the testing data). All *k* joint probabilities are multiplied together for a final score. In practice, we instead work with the natural logarithm of the probabilities and add them together to prevent rounding errors. Adding logarithms of probabilities is mathematically equivalent to multiplying probabilities.

Cross-validation is a somewhat blunt tool – though it is a perfectly general way to compare any two models on any given dataset, it requires relatively large amounts of data to return meaningful results. We therefore combine results within participants and across sessions. We consider two models to have meaningfully different scores if their final joint probability differs by a factor of 100 (equal to a natural log difference of 4.61).

### Results and Discussion

**Figure A1.**
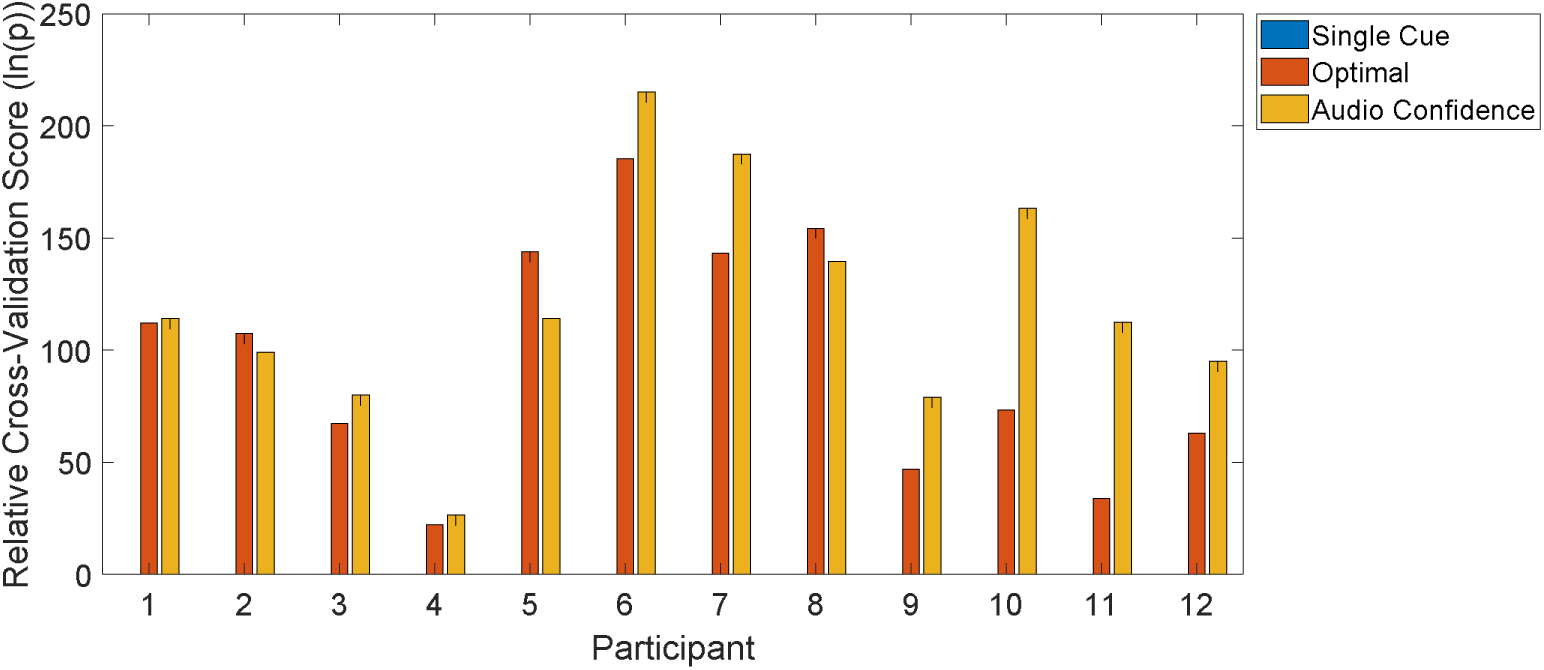
Relative cross-validation scores are the raw log probability (i.e. the final result of the cross-validation procedure) minus the lowest score. Higher scores are better. Scores are combined within participants but across sessions.

Results here are broadly consistent with the results in the main paper. For all 12 participants, the Single Cue model was meaningfully worse than the other two. The Optimal model was meaningfully better than the other two models for three participants (2, 5, and 8). One participant (1) had similar results for both the Optimal model and the Audio Confidence model. The remaining eight participants were meaningfully better fit with the Audio Confidence model than the other two. This suggests conceptually that all participants were combining the audio and visual cue. However, a minority were combining optimally and most were under-weighting the audio cue. This fits with the conclusions presented in the main text (i.e. as an average, the audio cue was underweighted, and the cues were not used to maximum efficiency).

In summary, the cross-validation analysis bolstered our confidence in the interpretation that is given in the main text.

## References

Abboud, S., Hanassy, S., Levy-Tzedek, S., Maidenbaum, S., & Amedi, A. (2014). EyeMusic: Introducing a “visual” colorful experience for the blind using auditory sensory substitution. Restorative Neurology and Neuroscience, 32(2), 247–257. https://doi.org/10.3233/RNN-130338

Alais, D., & Burr, D. (2004). The Ventriloquist Effect Results from Near-Optimal Bimodal Integration. Current Biology, 14(3), 257–262. https://doi.org/10.1016/j.cub.2004.01.029

Amedi, A., Hofstetter, S., Maidenbaum, S., & Heimler, B. (2017). Task Selectivity as a Comprehensive Principle for Brain Organization. In Trends in Cognitive Sciences (Vol. 21, Issue 5, pp. 307–310). Elsevier Ltd. https://doi.org/10.1016/j.tics.2017.03.007

Auvray, M., & Myin, E. (2009). Perception With Compensatory Devices: From Sensory Substitution to Sensorimotor Extension. Cognitive Science, 33, 1036–1058. https://doi.org/10.1111/j.1551-6709.2009.01040.x

Burr, D., & Gori, M. (2012). Multisensory Integration Develops Late in Humans. In The Neural Bases of Multisensory Processes. CRC Press/Taylor & Francis. http://www.ncbi.nlm.nih.gov/pubmed/22593886

Chebat, D.-R., Maidenbaum, S., & Amedi, A. (2015). Navigation Using Sensory Substitution in Real and Virtual Mazes. PLOS ONE, 10(6), e0126307. https://doi.org/10.1371/journal.pone.0126307

Daee, P., Mirian, M. S., Ahmadabadi, M. N., Brenner, E., & Tenenbaum, J. (2014). Reward Maximization Justifies the Transition from Sensory Selection at Childhood to Sensory Integration at Adulthood. PLoS ONE, 9(7), e103143. https://doi.org/10.1371/journal.pone.0103143

Deroy, O., & Auvray, M. (2012). Reading the World through the Skin and Ears: A New Perspective on Sensory Substitution. Frontiers in Psychology, 3, 457. https://doi.org/10.3389/fpsyg.2012.00457

Ernst, M. O. (2006). A Bayesian view on multimodal cue integration. In G. MKnoblich, I. Thornton, M. Grosjean, & M. Shiffran (Eds.), Human body perception from the inside out: Advances in visual cognition (pp. 105–131). Oxford University Press.

Ernst, M. O., & Banks, M. S. (2002). Humans integrate visual and haptic information in a statistically optimal fashion. Nature, 415(6870), 429–433. https://doi.org/10.1038/415429a

Ernst, M. O., Rohde, M., & van Dam, L. C. J. (2016). Statistically Optimal Multisensory Cue Integration: A Practical Tutorial. Multisensory Research, 29(4–5), 279–317. https://doi.org/10.1163/22134808-00002510

Getty, D. J. (1975). Discrimination of short temporal intervals: A comparison of two models. Perception & Psychophysics, 18(1), 1–8. https://doi.org/10.3758/BF03199358

Goeke, C. M., Planera, S., Finger, H., & König, P. (2016). Bayesian Alternation during Tactile Augmentation. Frontiers in Behavioral Neuroscience, 10, 187. https://doi.org/10.3389/fnbeh.2016.00187

Hershenson, M. (1962). Reaction time as a measure of intersensory facilitation. Journal of Experimental Psychology, 63(3), 289–293. https://doi.org/10.1037/h0039516

Hillis, J. H., Ernst, M. O., Banks, M. S., & Landy, M. S. (2002). Combining sensory information: Mandatory fusion within, but not between, senses. Science, 298(5598), 1627–1630. https://doi.org/10.1126/science.1075396

Knill, D. C., & Pouget, A. (2004). The Bayesian brain: the role of uncertainty in neural coding and computation. Trends in Neurosciences, 27(12), 712–719. https://doi.org/10.1016/j.tins.2004.10.007

Kolarik, A. J., Cirstea, S., Pardhan, S., & Moore, B. C. J. (2014). A summary of research investigating echolocation abilities of blind and sighted humans. Hearing Research, 310, 60–68. https://doi.org/10.1016/j.heares.2014.01.010

König, S. U., Schumann, F., Keyser, J., Goeke, C., Krause, C., Wache, S., Lytochkin, A., Ebert, M., Brunsch, V., Wahn, B., Kaspar, K., Nagel, S. K., Meilinger, T., Bülthoff, H., Wolbers, T., Büchel, C., & König, P. (2016). Learning new sensorimotor contingencies: Effects of long-term use of sensory augmentation on the brain and conscious perception. PLoS ONE, 11(12). https://doi.org/10.1371/journal.pone.0166647

Macmillan, N. A., & Creelman, C. D. (2004). Detection Theory: A User’s Guide: 2nd edition. In Detection Theory: A User’s Guide: 2nd edition. Psychology Press. https://doi.org/10.4324/9781410611147

Maidenbaum, S., & Abboud, S. (2014). Sensory substitution: Closing the gap between basic research and widespread practical visual rehabilitation. Neuroscience & Biobehavioral Reviews, 41, 3–15. https://doi.org/10.1016/J.NEUBIOREV.2013.11.007

Maidenbaum, S., Hanassy, S., Abboud, S., Buchs, G., Chebat, D.-R., Levy-Tzedek, S., & Amedi, A. (2014). The EyeCane, a new electronic travel aid for the blind: Technology, behavior & swift learning. Restorative Neurology and Neuroscience, 32(6), 813–824. https://doi.org/10.3233/RNN-130351

Maloney, L. T., & Mamassian, P. (2009). Bayesian decision theory as a model of human visual perception: Testing Bayesian transfer. Visual Neuroscience, 26(01), 147. https://doi.org/10.1017/S0952523808080905

Miller, J. (1982). Divided attention: Evidence for coactivation with redundant signals. Cognitive Psychology, 14(2), 247–279. https://doi.org/10.1016/0010-0285(82)90010-X

Moors, A., & De Houwer, J. (2006). Automaticity: A theoretical and conceptual analysis. Psychological Bulletin, 132(2), 297–326. https://doi.org/10.1037/0033-2909.132.2.297

National Public Radio. (2011). Blindness No Obstacle To Those With Sharp Ears. https://www.npr.org/2011/03/13/134425825/human-echolocation-using-sound-to-see?t=1585756613317

Negen, J., Wen, L., Thaler, L., & Nardini, M. (2018). Bayes-Like Integration of a New Sensory Skill with Vision. Scientific Reports, 8(1), 16880. https://doi.org/10.1038/s41598-018-35046-7

Norman, L. J., & Thaler, L. (2019). Retinotopic-like maps of spatial sound in primary ‘visual’ cortex of blind human echolocators. Proceedings of the Royal Society B: Biological Sciences, 286(1912), 20191910. https://doi.org/10.1098/rspb.2019.1910

Otto, T., & Mamassian, P. (2017). Multisensory decisions: the test of a race model, its logic, and power. Multisensory Research, 30(1), 1–24.

Pouget, A., Beck, J. M., Ma, W. J., & Latham, P. E. (2013). Probabilistic brains: knowns and unknowns. Nature Neuroscience, 16(9), 1170–1178. https://doi.org/10.1038/nn.3495

Rahnev, D., & Denison, R. N. (2018). Suboptimality in perceptual decision making. Behavioral and Brain Sciences, 41, e223. https://doi.org/10.1017/S0140525X18000936

Shams, L., Kamitani, Y., & Shimojo, S. (2002). Visual illusion induced by sound. Cognitive Brain Research, 14(1), 147–152. https://doi.org/10.1016/S0926-6410(02)00069-1

Striem-Amit, E., Cohen, L., Dehaene, S., & Amedi, A. (2012). Reading with Sounds: Sensory Substitution Selectively Activates the Visual Word Form Area in the Blind. Neuron, 76(3), 640–652. https://doi.org/10.1016/j.neuron.2012.08.026

Thaler, L., De Vos, H. P. J. C., Kish, D., Antoniou, M., Baker, C. J., & Hornikx, M. C. J. (2019). Human Click-Based Echolocation of Distance: Superfine Acuity and Dynamic Clicking Behaviour. JARO - Journal of the Association for Research in Otolaryngology, 20(5), 499–510. https://doi.org/10.1007/s10162-019-00728-0

Thaler, L., & Goodale, M. A. (2016). Echolocation in humans: an overview. Wiley Interdisciplinary Reviews: Cognitive Science, 7(6), 382–393. https://doi.org/10.1002/wcs.1408

Zhang, X., Reich, G. M., Antoniou, M., Cherniakov, M., Baker, C. J., Thaler, L., Kish, D., & Smith, G. E. (2017). Human echolocation: waveform analysis of tongue clicks. Electronics Letters, 53(9), 580–582. https://doi.org/10.1049/el.2017.0454

